# Grey mouse lemurs, *Microcebus murinus,* are a relevant model to study gut microbiome flexibility in response to diet changes

**DOI:** 10.64898/2026.08.17.744835

**Authors:** Maxime Naour, Isabelle Grit, Patricia Parnet, Hervé M. Blottière, Jérémy Terrien

## Abstract

The gut microbiota is a key player in energy balance, impacting both digestion efficiency and the production of metabolites involved in metabolism. Its composition is highly adaptable, especially in response to diet. Changes in human diet and lifestyle over time – from active, fibre-rich diets to sedentary habits with calorie-dense foods – have likely contributed to the rise in metabolic diseases. Rodent models are widely used to study the links between diet, microbiota and metabolism. However, they have important limitations (e.g. artificial environments, uniform diets and biological differences from humans) which can affect the translation of findings to humans. While mice and humans differ in their microbiota species, they do share some functional similarities. The grey mouse lemur (*Microcebus murinus*) has been proposed as a promising alternative model. This small primate experiences strong seasonal changes in food availability, leading to distinct physiological states (energy-saving in winter vs active in summer), even in captivity. It is increasingly recognized as a valuable model for biomedical research, supported by recent genomic and molecular advances. However, its gut microbiota has not yet been the subject of study. Consequently, the present study focuses on investigating the gut microbiota of the grey mouse lemur, with a particular emphasis on how these microbiota vary under different dietary regimens. The microbiota of animals fed the standard colony diet was dominated by *Prevotella*, *Bifidobacterium, Megamonas*, *Streptococcus*, *Megasphaera* and *Lactococcus*, showing an *Prevotella* driven enterosignature. We showed that switch from a classical control diet to 3 different diets resulted in change on microbiota composition that is associated with functional redundancy. The present work underline the interest of *Microcebus murinus* as model for diet and lifestyle studies in relationship with metabolic diseases.

## Introduction

The gut microbiota is at the interface between environmental food availability and the organism and has been shown to play a key role in energy homeostasis [Wang et al, 2017]. Specifically, the gut microbiota has major impact on the efficiency of digestion and the metabolites that are made available for energy metabolism [Riedl et al, 2017; Scheithauer et al, 2016]. It is well described that gut microbiota composition is flexible, particularly in response to food composition [Doré & Blottière, 2015; Kaczmarek et al; 2017; Hauffe & Barelli, 2019; Carmody et al, 2019; Amato et al, 2015]. The modification of diet composition and feeding habits during human evolution – i.e. switching from physically active hunter– gatherers with a diet rich in fibers but low in sugar to sedentary persons with low physical activity and easy access to extremely rich food – may also have contributed to promote energy imbalance and the development of metabolic diseases [Carter et al, 2023; O’Keefe 2019].

Following the pioneer work from Jeffrey Gordon’s lab [Bäckhed et al, 2004; Bäckhed et al, 2005; Turnbaugh et al, 2006], many studies have been using rodent models to study mechanisms linking diet, microbiota and metabolic disorders. However, these rodent models have advantages but also major limitations that have been discussed [Nguyen et al, 2015]. Among the limitation, one can discuss the housing condition (specific pathogen free animal facilities), the standardized chow diet (rodent eat all lifelong the same food), coprophagy, genetic homogeneity (inbred mouse strains), difference in anatomy, genetics and physiology. Of course, one has to critically review the translational perspective of the pathophysiological model selected [Preguiça et al, 2020]. Regarding the microbiome, although major differences exist at the taxonomical level between mouse and human, indeed few species are shared between the two, at the functional level closer similarities have been observed [Xiao et al, 2015; Beresford-Jones et al, 2022].

The grey mouse lemur (*Microcebus murinus*) as an animal model, is an omnivorous primate that is very relevant in many aspects to address the questions. Indeed, this small-sized (body weight ∼100g), nocturnal species originates from Madagascar, which environmental constraints have shaped organisms with very specific life-history traits. Indeed, animals have to face two major seasons with very different energy constraints: a cold and dry winter (short days) with very low (if none) food availability; a hot and wet summer (long days) with high food availability [Wright, 1999]. Mouse lemurs are opportunistic and can eat fruits, flowers, insect exudates, arthropods and plant gums, and sometimes small vertebrates and leaves, depending on their seasonal availability [Radespiel et al, 2006; Dammhahn et al, 2008]. The environmental constraints have driven adaptive responses leading to very contrasted physiological and behavioral states between seasons, *i.e.* a hypometabolic and reproductively quiescent state during winter [Schmid & Speakman, 2020] vs a very active and reproductive state during summer [Rina Evasoa et al, 2018]. Such seasonality is strikingly maintained in captivity, while environmental conditions (including food availability and ambient temperature) are maintained high and constant, by the only manipulation of day-length (6 months of short days followed by 6 months of summer) [Perret & Aujard, 2001]. Mouse lemurs are internationally recognized as highly relevant for investigations in primate biology and behavior, with a specific emphasis on biomedical studies and recently highlighted as an emerging model organism [Camp, 2025]. More light has recently being shed on this model as a new assembly of the genome has been recently published [Ezran et al, 2025], together with a detailed single cell transcriptomic atlas from 27 organs providing all the molecular tools to perform high throughput analyses [Tabula Microcebus consortium, 2025]. However, to the best of our knowledge, the gut microbiota of the grey mouse lemur has never been analyzed, either in its natural habitats of Madagascar or in captivity in the laboratory. Though, the previously reported microbiota analysis from gray-brown mouse lemurs (*M. griseorufus*), a closely related species, gave light on the lemurs microbiome [Wasimuddin et al, 2022]. This work exemplified the consequences of human-driven habitat disturbance, which results in diminished gut bacterial diversity and subsequent alterations in microbial metabolic functions. This underscores the notion that the gut microbiome can serve as a biological indicator of environmental or nutritional stressors and host health. It highlights the need to include microbiome analysis in study using this mighty model organism.

In the present work, we report on the study of the gut microbiota of *M. murinus* in captivity and on microbiota modulation by different diets.

## Methods

### Experimental design and sample collection

#### Animal husbandry

Grey mouse lemurs (*M. murinus*) were born in the captive population at Brunoy (MNHN, France, approval license no. F91-114-1) and raised in optimal conditions to ensure their survival, reproduction, growth, and well-being. Water and food are available *ad libitum* throughout the year and the animals are kept in same-sex groups of one to six animals per cage. The breeding rooms are maintained at a constant, controlled temperature and humidity (23 to 26°C and 55% on average, respectively). Seasonal entrainment is maintained under captive conditions through changes in daylength (6 months of short days followed by 6 months of long days) therefore allowing physiological and behavioral modifications throughout the year. During the 6-months period of long days, light is maintained artificially from 3 a.m. to 5 p.m. (i.e., 14 hours of light per day), while lights are kept on artificially from 7 a.m. to 5 p.m. during short days (i.e., 10 hours of light per day).

#### Experimental procedures

All experimental procedures were approved by the Animal Welfare Council of UMR 7179, the Cuvier Ethics Committee for the Care and Use of Experimental Animals at the Muséum national d’Histoire Naturelle, and authorized by the Ministry of Higher Education, Research, and Innovation (No. APAFIS#27250-2020091510519374 v3) and complied with European ethical regulations for the use of animals in biomedical research. Eleven adult male mouse lemurs (2.9 +/- 0.2 years old; mean body mass 85.0 +/- 4.7 g) were followed in a within-animal cross-over between February and April 2021. Each animal first received the standard colony diet, a control mash (CTL) reproducing its everyday ration, and was then challenged for three days with a single alternative preparation of matched daily energy (about 25 kcal per day): either raw meat (RAW), the same meat once cooked (COOK), or a fat-enriched mash (HF), whose detailed formulations are given in Table 1. After the high-fat challenge, animals were returned to the control diet and sampled again, so that their recovery could be followed as a distinct condition (CTL post-HF). Stools were collected non-invasively throughout each animal’s series, spanning baseline, challenge and, where relevant, recovery; in total, 31 fecal samples were obtained from the eleven animals, comprising 13 taken at baseline and 5 after recovery on the control diet, together with 4 under the high-fat, 4 under the raw and 5 under the cooked diet (see Table 2). Only males were studied, to avoid the pronounced sex-specific reproductive and body-mass seasonality of this species; because sampling spanned the transition between photoperiodic seasons, season was retained as a covariate in sensitivity analyses. An extraction blank and a positive control were processed and sequenced alongside the biological samples to monitor reagent contamination and to verify the sequencing and bioinformatic workflow. Diet effects were examined both against the control, by contrasting the high-fat and the raw diets with it, and within animals on paired samples, by comparing the recovery state with the high-fat challenge and with the baseline, and the cooked diet with the raw one.

**Table 2:** Animal id, date of sampling, diet at time of sampling and, in parenthesis, sequencing sample number.

|  | D0 | D6 | D12 | D21 | D0 | D3 | D6 |
| --- | --- | --- | --- | --- | --- | --- | --- |
| *Date*<br>ID# | *Febr 12* | *Febr 18* | *March 2* | *March 11* | *Apr 2* | *Apr 5* | *Apr 8* |
| 297CBB | CTR (9) |  | CTR (14) | CTR (16) |  |  |  |
| 374B | CTR (29) |  |  | CTR (28) |  |  |  |
| 374A | CTR (1) | HFD (6) | CTR (8) | CTR (18) |  |  |  |
| 275BBE | CTR (13) | HFD (22) |  |  |  |  |  |
| 283BCB | CTR (31) | HFD (2) | CTR (27) |  |  |  |  |
| 289BCB |  | HFD (12) | CTR (23) | CTR (21) |  |  |  |
| 225BCA |  |  |  |  | CTR (25) | RAW (20) | COOK (4) |
| 262DBA |  |  |  |  | CTR (7) | RAW (5) | COOK (17) |
| 294AF |  |  |  |  | CTR (32) | RAW (19) | COOK (3) |
| 314BBA |  |  |  |  | CTR (15) |  | COOK (11) |
| 364AAA |  |  |  |  | CTR (24) | RAW (26) | COOK (10) |

Sequencing samples number 30 and 33 were blank and positive control, respectively.

#### Host phenotyping

Alongside each stool collection, animals were weighed and phenotyped for body composition and blood biochemistry. Body mass (BM) was recorded in grams. Body composition was measured by and expressed as the percentage of body mass accounted for by fat (%Fat), by free body water (%Fluid) and by lean tissue (%Lbm). Blood was collected and analyzed for glucose (Gly), total cholesterol (Chl), high-density-lipoprotein cholesterol (HDL) and triglycerides (TRI), all expressed in mg/dL. Low-density-lipoprotein cholesterol and the total-cholesterol-to-HDL ratio were derived from these measurements rather than assayed. Body composition was obtained for all 31 samples; the blood panel was available for 21 to 22 of them and was not collected under the raw diet. Total cholesterol reached the upper limit of the assay (500 mg/dL) in 10 of the 21 measured samples and is therefore right-censored above that value.

## 16S rRNA Gene Amplicon Sequencing

Total DNA from all stool samples was extracted using the QIAamp® Fast DNA Stool Mini Kit (Qiagen) and then quality-checked by NanoDrop (260/280 > 1.5; 260/230 > 1.2), Qubit dsDNA BR assay (target ≈ 50 ng/µL) and Agilent 2100 TapeStation (DIN > 5.5). DNA samples were normalized to 5 ng/µL (5 µL DNA + 45 µL nuclease-free water), arrayed in a 96-well plate, and then amplified in two PCR steps targeting the V3-V4 region of the 16S rRNA gene.

In the first amplification (PCR1), overhang primers C1_ovh-ilm_V3-V4_341F (5ʹ-TCGTCGGCAGCGTCAGATGTGTATAAGAGACAG CCTACGGGNGGCWGCAG-3ʹ) and C1_ovh-ilm_V3-V4_804R5ʹ-GTCTCGTGGGCTCGGAGATGTGTATAAGAGACAG GACTACHVGGGTATCTAATC C-3ʹ) were used under the following thermal profile: 95 °C for 3 min; 16 cycles of 95 °C for 30 s, 55 °C for 30 s and 72 °C for 30 s; and a final 5 min extension at 72 °C. PCR1 products were purified with 0.8× AMPure XP beads. With the same temperature settings, the purified amplicons then entered indexing PCR (PCR2) with Nextera XT v2 dual-index primers (i7 indices N718-N728, i5 indices S502-S511) for 9 cycles, followed by clean-up with 1.0× AMPure XP beads.

Indexed libraries were size-validated on an Agilent 2100 TapeStation (DNA 1000 Kit), with major peaks between 450-700 bp (and no detectable profile for extraction blanks). They were then quantified on a Qubit Flex fluorometer using the 1× dsDNA High Sensitivity Assay Kit (Thermo Fisher), converted to nanomolarity based on an average fragment size of 602 bp, and only those adjusted to > 4 nM (blanks < 1 nM) were retained. Each library (100-190 nM) was diluted 1:10 in 10 mM Tris pH 8.5 (to ∼10- 19 nM), and 2 µL of each dilution were pooled equimolarly. The pooled libraries were denatured with 0.2 N NaOH, spiked with ∼22 % PhiX control, and sequenced as 250 bp paired-end reads at 4 pM on an Illumina MiSeq instrument using the MiSeq V2 500-cycle SBS kit (250-8-8-250 run). Raw BCL files were converted on-instrument to per-sample FASTQ files for downstream analysis.

### Pre-processing of 16S rRNA metabarcoding data

Raw 2×250 bp paired-end MiSeq FASTQ files were checked with FastQC v0.11.9 and merged by MultiQC v1.12 [Andrew, 2010; Ewels et al, 2016], then pre-processed in FROGS v5.0.0 [Escudié et al, 2018] using the Illumina denoising pipeline. Read pairs were merged using PEAR [Zhang et al, 2014], enforcing a minimum overlap of 20 bp and allowing up to 15 % mismatches. Primer sequences (forward: 5ʹ-CCTACGGGNGGCWGCAG-3ʹ; reverse: 5ʹ-GGATTAGATACCCBDGTAGTC-3ʹ) were then trimmed, and any reads missing either primer were discarded. Merged sequences shorter than 200 bp, longer than 490 bp, or containing ambiguous bases (N) were removed. The remaining high-quality reads were dereplicated and denoised using the DADA2 algorithm [Callahan et al, 2016] in pseudo-pooling mode – specifying 250 bp read lengths in both directions, a 0.15 maximum mismatch rate and initial amplicon-size bounds of 200-490 bp – which were then narrowed to 390-470 bp based on the diagnostic report. Chimeric sequences were identified de novo and removed with UCHIME [Edgar et a, 2011]. Finally, abundance- and prevalence-based filtering (retaining ASVs with ≥ 0.005 % relative abundance in more than one sample and excluding phiX contaminants) produced the final filtered ASV dataset for downstream taxonomic affiliation.

### Taxonomic assignment and functional potential inference

Taxonomic affiliation of the filtered ASV was processed in FROGS v5.0.0 using the GTDB_08-RS214 16S-ITS-23S reference database (doi:10.57745/APDPLQ). Assignment was performed with the taxonomic_affiliation.py module employing the RDP classifier algorithm [Wang et al, 2007]. The resulting BIOM table was converted to tabular form to identify ASVs with multiple candidate affiliations, which were then ranked by BLAST percent-subject coverage and manually curated. A newick-formatted phylogenetic tree of ASVs was subsequently inferred with the tree.py module.

Inference of community functional potential was performed using the FROGSFUNC v5.0 pipeline. Initially, ASVs were aligned to reference HMM profiles with HMMER [Eddy, 2011] and placed into the PICRUSt2 [Douglas et al, 2020] backbone phylogeny of 20,000 prokaryotic 16S rRNA sequences using EPA-NG [Barbera et al, 2019]. Placement results were integrated into the reference phylogeny using GAPPA [Czech et al, 2020]. NSTI scores were computed for each ASV to quantify its phylogenetic distance to the nearest sequenced genome. ASVs with NSTI values above 0.64 (the 95th percentile) were excluded. Subsequently, maximum-parsimony hidden-state reconstruction in castor-R [Louca & Doebeli, 2018] was used to predict per-ASV abundances of key functional categories (EC, KO, COG, PFAM, TIGRFAM, PHENO); these predicted counts were then normalized by the estimated 16S copy number and ASVs exceeding the NSTI threshold were removed. Finally, predicted gene families were mapped to MetaCyc [Caspi et al, 2016; 2020] and KEGG reactions [Kanehisa et al,2025] and pathways using MinPath [Ye & Doak, 2009], with both copy-number–normalized and raw analyses for EC and KO annotations.

### Statistical analysis

Data processing and community profiling. All downstream analyses were conducted in R (v4.5.3) [R Core Team, 2024], with three phyloseq [McMurdie & Holmes, 2013] objects supporting taxonomic community profiling and functional potential inference (KEGG-pathways levels and EC-genes levels). Low-abundance ASVs were removed by retaining only those with at least 10 counts in at least 10 % of the samples, reducing the dataset from 787 to 713 ASVs, and taxonomic analyses were carried out at genus and species (named-ASV) resolution. Rarefaction curves and per-group sequencing depth were inspected to confirm adequate and comparable coverage. Depending on the analysis, counts were used after total-sum scaling for relative-abundance displays or after a robust centered-log-ratio transform [Gloor et al, 2017; Martino et al, 2019] for compositional analyses, so that samples were compared as compositions rather than as raw proportions. Community composition was displayed as per-sample stacked bar plots at genus and species level, as phylum-to-family alluvial diagrams and as core- microbiota plots relating each taxon’s prevalence to its mean abundance; the sharing of predicted functions among taxa (functional redundancy) was summarized as the effective number of genus carriers per KEGG pathway.

#### Alpha diversity

Within-sample diversity was summarized by observed richness and Pielou’s evenness. [Pielou, 1966]. Because the design is longitudinal, differences between diets were tested with mixed models comprising a diet fixed effect and a random intercept for animal identity, using generalized linear mixed models for count-type richness [Bates et al, 2015; Brooks et al, 2017] and linear or beta mixed models for bounded evenness [Ferrari & Cribari-Neto, 2004]; pairwise diet differences were obtained from estimated marginal means [Lenth, 2025]. Diversity was analyzed at the species, KEGG-pathway and EC-gene levels.

#### Beta diversity

Between-sample dissimilarity was computed as the Aitchison distance on centered-log- ratio-transformed abundances and summarized by principal component analysis. Separation between diets was tested by permutational multivariate analysis of variance (PERMANOVA) [Anderson, 2001; Oksanen et al, 2025] on the Aitchison distance [Aitchison, 1982], and because a significant test may reflect a shift in community location or a difference in dispersion, every PERMANOVA was paired with a test of multivariate homogeneity of group dispersions (betadisper) [Anderson, 2006]; a location effect was inferred only when the dispersion test was non-significant. Beta diversity was analyzed at the species and KEGG-pathway levels.

#### Differential abundance

Differentially abundant features were identified on centred-log-ratio- transformed counts with LinDA [Zhou et al, 2022]. For each feature the effect size is the log2 fold-change of the test diet relative to its reference, and significance was controlled by the Benjamini-Hochberg false-discovery rate [Benjamini & Hochberg, 1995] within each contrast. The five contrasts (HF vs CTL, CTL post-HF vs HF, CTL post-HF vs CTL, RAW vs CTL and COOK vs RAW) were analysed at the species, KEGG-pathway and EC-gene levels, with the within-animal contrasts modelled on paired samples.

#### Supervised discrimination

Sparse partial-least-squares discriminant analyses (sPLS-DA) [Lê Cao et al, 2011] were fitted with the mixOmics package [Rohart et al, 2017] to identify the features that best separate each pair of diets, ranking their contributions by Variable Importance in Projection (VIP); model performance and feature stability were assessed by nested leave-one-animal-out cross- validation. sPLS-DA was applied to species and KEGG pathways for the five contrasts.

#### Enterosignatures

Enterosignatures [Frioux et al, 2023] were derived by non-negative matrix factorization with a Kullback-Leibler objective [Lee & Seung, 1999], which expresses each sample as an additive mixture of a few signatures rather than assigning it to a single type. The number of signatures was chosen per data scope by Owen-Perry repeated bi-cross-validation [Owen & Perry, 2009] with a one-standard-error parsimony rule and verified by leave-one-animal-out and animal-level bootstrap resampling; at the sample sizes available here the decision was restricted to the reliably distinguishable choice between a single continuous gradient and a minimal two-state structure. Signatures were named after their co-dominant driver features and visualized with PHATE [Moon et al, 2019], a non-linear embedding of the compositional geometry that preserves continuous transitions between samples.. Enterosignatures were computed at the genus, species, EC-gene and KEGG-pathway levels.

#### Co-occurrence networks

Feature co-occurrence was described with LUPINE partial-correlation networks [Kodikara & Lê Cao, 2025], in which an edge represents a conditional association between two features after accounting for the others. An edge was retained only when its permutation p-value passed a cutoff and the magnitude of the partial correlation exceeded a floor calibrated per layer and per cohort, and edge color and opacity encoded the sign and strength of the association. Networks were partitioned into guilds with the Louvain algorithm [Blondel et al, 2008] and candidate keystone features were flagged from their Guimerà-Amaral within-module and among-module connectivity (Zi- Pi) roles, with nodes colored and grouped by phylum [Guimerà & Amaral, 2005]. Networks were built for the all-diet and control-only cohorts and one network was drawn per enterosignature at the species and KEGG-pathway levels. Because associations at these sample sizes are hypotheses rather than validated interactions, the network and keystone results are treated as exploratory.

#### Microbiome-phenotype associations

Associations between microbiome features and host phenotypes were quantified by Spearman rank correlation (ρ) [Spearman, 1904] between centered-log- ratio-transformed abundances, obtained after Bayesian multiplicative replacement of zeros [Palarea- Albaladejo & Martín-Fernández, 2015], and each phenotype. Correlations were computed separately for each of the five diet contrasts on the pooled samples of both arms, so that a coefficient describes how a feature tracks a phenotype across the dietary transition; it therefore incorporates the shift between the two diets by construction. A repeated-measures formulation was considered but not adopted: with 1.5 to 2.3 samples per animal, a random intercept for animal identity is not identifiable, so the model would collapse to an ordinary regression while appearing to account for the dependence [Bakdash & Marusich, 2017]; rank correlation with leave-one-animal-out resampling was preferred as a more robust and more transparent description. Three admissibility rules were applied before a coefficient was reported: a feature had to be detected in at least half of the animals of both arms, failing which ρ would express a presence-absence contrast rather than an association; both arms had to contribute at least one sample with a measured phenotype, failing which the coefficient would be a within-group correlation carrying a contrast label; and at least six samples from four animals were required. Sign stability was assessed by refitting every correlation with one animal removed at a time. P-values were adjusted by the Benjamini-Hochberg false-discovery rate [Benjamini & Hochberg, 1995] within each contrast-by-phenotype family, and the five features with the largest absolute correlation were retained per contrast and per phenotype, ties being broken by leave-one-animal-out sign stability. Derived phenotypes were excluded, low-density-lipoprotein cholesterol and the total-cholesterol-to-HDL ratio being exact functions of the measured panel (ρ = 0.90 between total and low-density-lipoprotein cholesterol), as was age, whose within-contrast increment is identical for every animal and therefore carries no variance. For display, the retained features were partitioned by consensus k-means over 1000 restarts, the pairwise co-clustering matrix being cut by average linkage and the number of clusters selected by the mean silhouette width [Rousseeuw, 1987; Monti et al, 2003]; heatmaps were drawn with ComplexHeatmap [Gu et al, 2016]. Associations were computed at the species, KEGG-pathway, EC-gene and KO-gene levels. Because the available sample sizes cannot support confirmatory inference and the p-values do not model the repeated measurements, these associations are reported as exploratory.

#### Statistical reporting and reproducibility

Analyses were run in R (v4.5.3) using, among other packages, phyloseq, vegan, ALDEx2, LinDA, mixOmics, lme4 and glmmTMB with emmeans, and igraph [Csárdi et al, 2006] and ggraph [Pedersen, 2024]. Multiplicity was controlled by the Benjamini-Hochberg false-discovery rate within each analysis, and findings are reported with effect sizes and measures of uncertainty rather than with p-values alone. The analyses of diet effects on diversity, composition and predicted function were pre-specified, whereas the enterosignature, co-occurrence-network and keystone analyses are exploratory. A fixed random seed of 42 was set for every stochastic step. The analysis code is available at https://github.com/MaximeNaour/, the raw 16S rRNA gene reads are deposited in the NCBI Sequence Read Archive (BioProject PRJNAXXXXXX), and sample metadata are provided in MIxS-compliant form as supplementary tables.

## Results

### A *Prevotella*-dominated reference community in the control-diet gut microbiome

In our study, all 11 male young adult *M. murinus* were fed the classical control diet, some of them have been stool sampled several times (Table 2). Gut microbiota composition was assessed using 16S rRNA gene sequencing. After sequencing, a minimum of 140 000 reads were obtained, and for all samples it was above the rarefaction threshold. Gut microbiota analysis of the samples from control group revealed an homogeneous composition a Bacteroidota-/*Prevotella*-dominated community that behaves as a single continuous enterosignature rather than discrete enterotypes. At the phylum level, Bacteroidota (previously Bacteroidetes) accounted for 33.0 %), and Bacillota (previously Firmicutes) accounted for 47.5% separated into Bacillota_A, 19.5%, Bacillota 15.6% and Bacillota_C, 12.4% constituted the most prominent taxonomic groups. Finally, Actinomycetota, and Actinomycetota (previously Actinobacteria) represented 14.8% and Pseudomonata (Proteobacteria) being relatively rare, only 2.2% (Figure 1A). At the family level, the microbiota of animal under control diet was dominated by *Bacteroidaceae*, *Lachnospiraceae*, *Bifidobacteriaceae*, *Streptococcaceae*, Selenomonadaceae, *Megasphaeraceae* and *Acutalibacteraceae*, followed by other minor family (Figure 1A). Figure 1B revealed substantial inter-individual and intra-individual heterogeneity variations at the genus level. *Prevotella* 19.4% leads, followed by *Bifidobacterium* 11.2%, *Megamonas* 7.3%, *Streptococcus* 6.7%, *Megasphaera* 4.3%, *Lactococcus* 4.2% Figure 1C illustrate the variation between samples at the genus level and highlight 68 genera that are highly abundant and very prevalent (> 90%) including *Prevotella*, *Bifidobacterium*, *Megamonas*, *Megasphaera*, *Ruminococcus*, *Phocaeicola* and *Bacteroides*. At the species level, *Prevotella copri B* represented 12.4% of all the species present, *Megamonas funiformis* 7.3%, *Streptococcus pasteurianus* 5.5% and *Bifidobacterium pseudocatenulatum* 4.3% that lead a diverse, individual-specific community (Fig. 1B, S1). Interestingly, figure 1D illustrate the functional redundancy between the main genera and the main KEGG pathways that were derived from the same 16S-based inference. These links described the attribution of predicted functional potential but did not constitute metabolomic validation. Community typing resolves to a single species-level enterosignature (K = 1), ES-*P. copri* (DMM Laplace and BIC both K = 1 for the CTL-only scope); the genus level likewise resolves to a single gradient (Fig. S1A) and the predicted-function level to a single signature ES-ko01051 (Fig. 1E, 1F). Per-signature LUPINE networks with Louvain guilds and Zi-Pi candidate keystones map the community’s organization at the species and KEGG-pathway levels (Fig. S1B, S1C).

**Figure 1.**
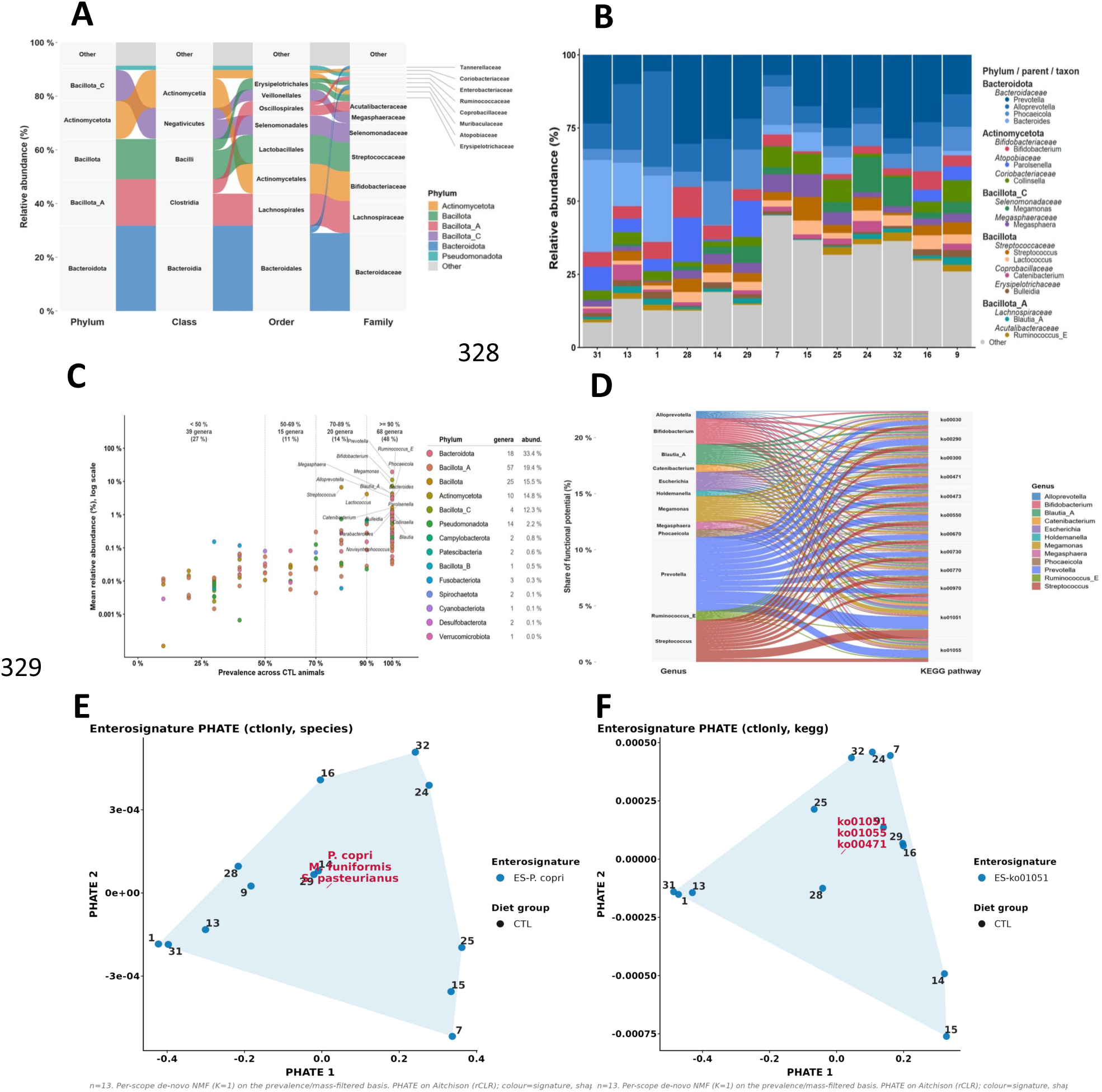
Characterization of the gut microbiome of captive-bred Microcebus murinus on the control diet. A) Taxonomic alluvial diagram tracing the flow of relative abundance from phylum to family, summarizing the dominant lineages of the control-diet microbiome; B) Per-sample stacked bar plot of genus-level relative abundance across the control-diet samples; the most abundant genera are color-coded and the remainder pooled as others; C) Core-microbiota plot: each taxon is positioned by its prevalence across animals against its mean relative abundance, defining the genera that consistently make up the control community; D) Functional-redundancy alluvial linking the main genera to the main predicted KEGG pathways, showing that several taxa carry the same functions; E) Enterosignature structure at the species level, shown as a PHATE embedding of the robust-CLR geometry; the community resolves into a single signature, ES-*P. copri*, i.e. a continuous *Prevotella*-dominated gradient rather than discrete types; F) Enterosignature structure at the KEGG-pathway level, likewise resolving to a single functional signature, ES-ko01051, consistent with the functional redundancy shown in D. Points are individual samples.

### A short dietary change reshapes taxonomic composition

Initially fed the control diet, some animal were put under a raw beef meat diet for 3 days before stool collection, followed by the same diet but with cooked beef meat. For some animal under control diet, a switch to high fat diet for 3 days was followed by a 12 and 21 days recovery before sampling. It is noteworthy that the 3 diet were isocaloric and, except of course for the HF diet, had the same balance between macronutrients, the only difference was from the ingredients used. For the HF diet, 60% were coming from lipids instead of 20% of calories in the control diet which was a high-carbohydrate diet. The analysis of the diets in a bomb calorimeter confirmed that all diets, as freshly prepared and given to the animals, were isocaloric, providing on average 1.02 Kcal/g of fresh mixture.

A three-day dietary change markedly reshuffles taxonomic composition in diet-specific directions; small-sample omnibus/per-feature tests rarely reach significance, so the diet fingerprints are read from group-mean composition and community typing. At the community level (beta diversity), there is a nominal diet signal at the genus level (PERMANOVA R² = 0.25, p = 0.018, q = 0.09), that is marginal at the species level (R² = 0.25, p = 0.059), both with significant dispersion heterogeneity (betadisper genus p = 0.017, species p = 0.004), = so not it is cleanly a location shift (Fig. 2A). However, analysis of the functional layers shows that there is no separation. (see Fig. 3).

**Figure 2.**
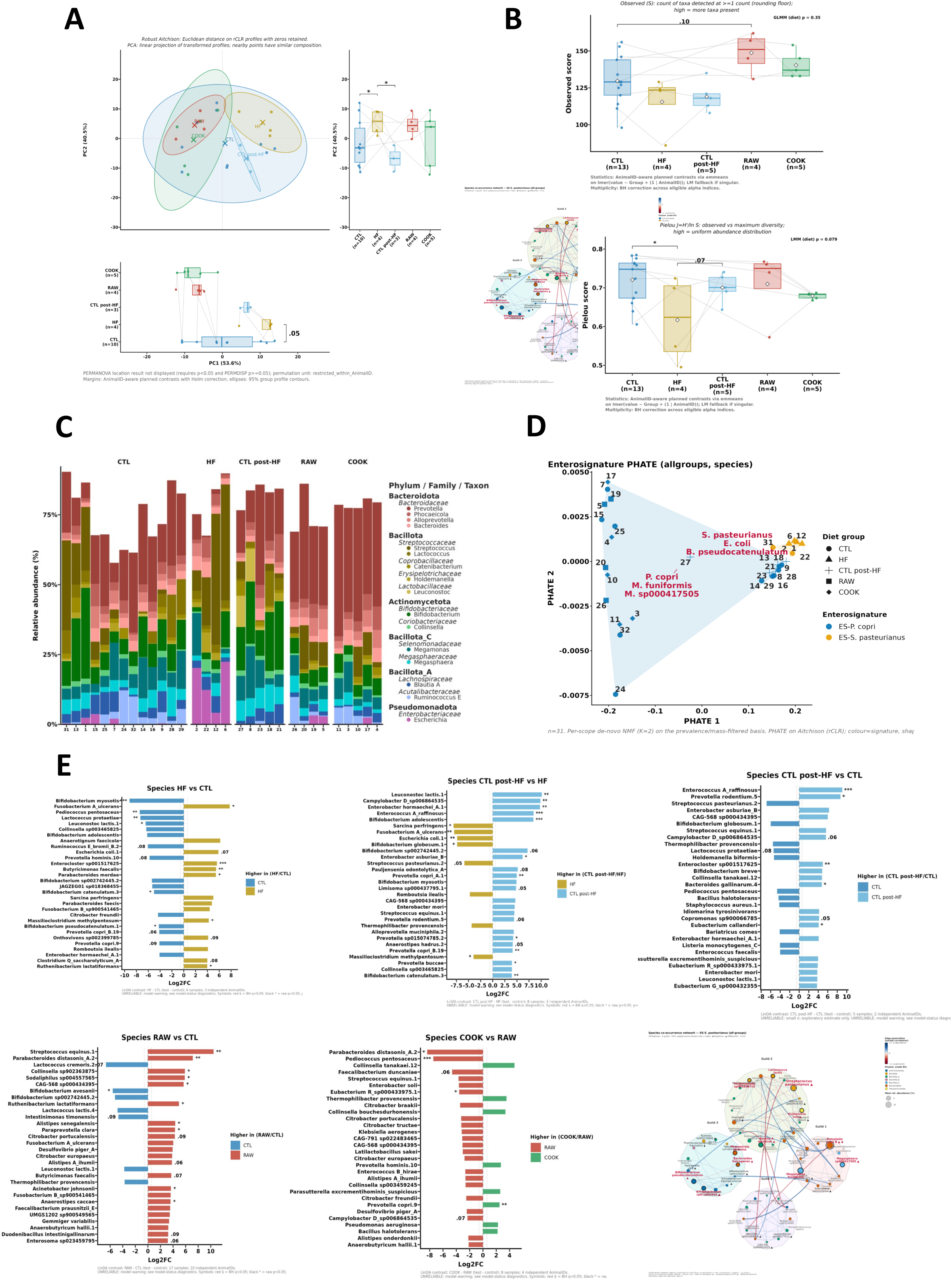
A short dietary change reshapes the taxonomic composition of the *Microcebus murinus* gut microbiome. A) Between-sample beta-diversity at the species level: PCA of all samples colored by diet, with axis marginals; group separation and dispersion were tested by PERMANOVA and betadisper on the Aitchison distance; B) Alpha diversity at the species level, observed richness (top) androbust-Aitchison Pielou evenness (bottom) by diet group, tested with a diet term in a mixed model with animal as a random effect; C) Per-sample stacked bar plot of genus-level relative abundance, samples grouped by diet, illustrating the compositional turnover; D) Enterosignature structure at the species level across all diets (PHATE embedding): two signatures now co-exist, ES-*P. copri* and ES-*S. pasteurianus*, with high-fat samples shifting toward the *Streptococcus*/ *Escherichia* signature. E) Species-level differential abundance (LinDA) for five diet contrasts, shown as signed effect-size bar plots (bars = log2 fold-change of the test diet relative to its reference, coloured by direction, FDR-controlled within each contrast), in the order HF vs CTL, CTL post-HF vs HF, CTL post-HF vs CTL, RAW vs CTL and COOK vs RAW. The co-occurrence network of the high-fat-associated signature ES-*S. pasteurianus* is shown alongside (conventions as in Fig. S1B).

**Figure 3.**
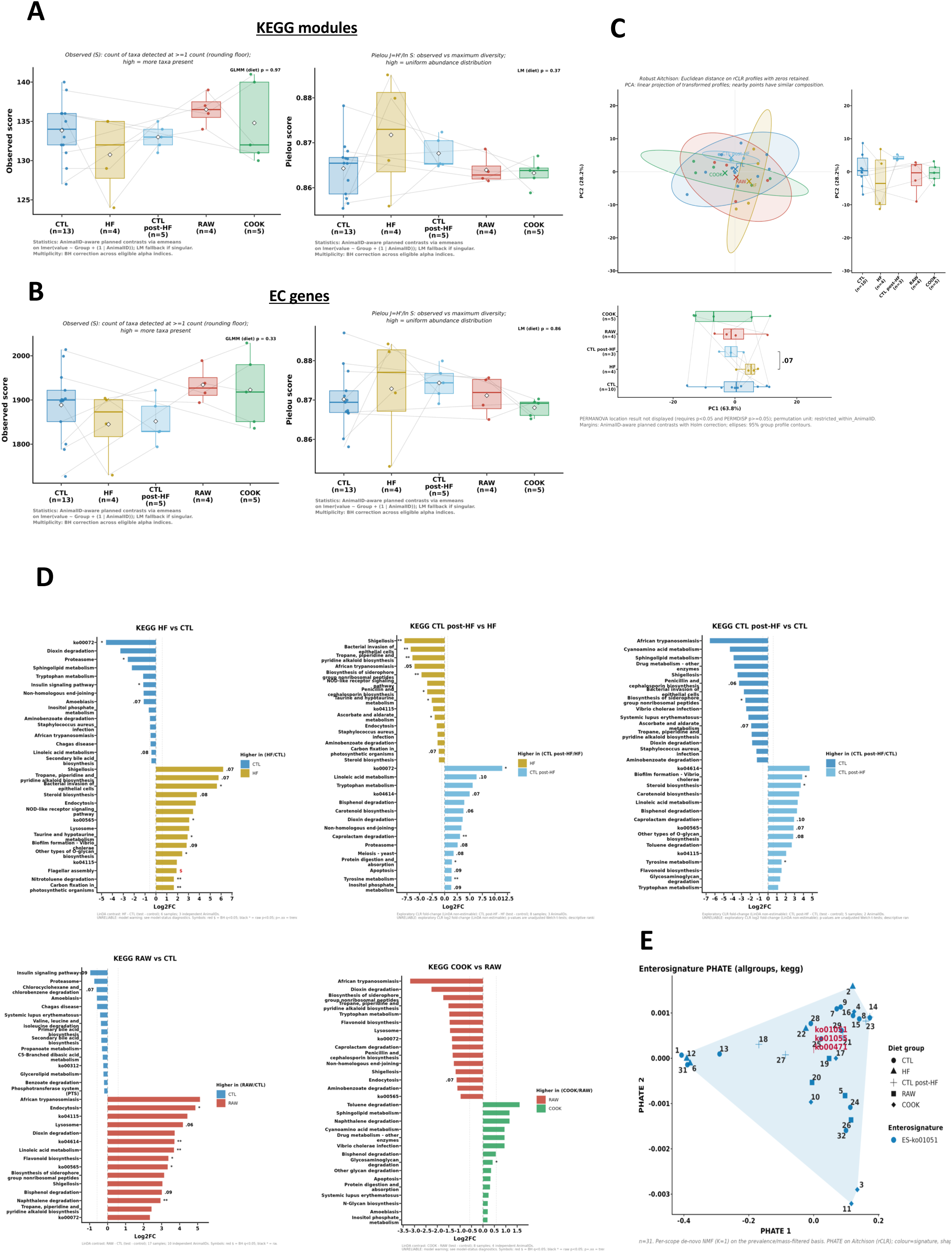
Dietary change has a comparatively modest effect on the predicted functional potential of the gut microbiome. A) Alpha-diversity at the KEGG-pathway level, observed richness (left) and Pielou evenness (right) by diet group (mixed model with animal as a random effect); B) Alpha-diversity at the EC-gene level, observed richness (left) and Pielou evenness (right) by diet group; C) Between-sample beta-diversity at the KEGG-pathway level: robust-Aitchison PCA of all samples with axis marginals (PERMANOVA/betadisper on the Aitchison distance); D) KEGG-pathway differential abundance (LinDA) for the five diet contrasts, shown as the top-15-per-direction signed bar plots (the fifteen most enriched pathways on each side; bars = log2 fold-change, FDR-controlled per contrast), in the same order as Figure 2E; E) Enterosignature structure at the KEGG-pathway level (PHATE embedding); the single functional signature underlines the functional redundancy of the community. Compared with the taxonomic level (Figure 2), the functional composition is markedly more stable across diets.

For the alpha diversity: no contrast survived Benjamini-Hochberg (BH) correction (minimum q ≈ 0.13). The interpretable trends are higher richness under raw (observed RAW vs CTL: species +21.5, p = 0.050; genus +15.1, p = 0.032) and lower evenness under high-fat (species Pielou HF vs CTL -0.104, p = 0.093; InvSimpson -8.70, p = 0.034) (Fig. 2B).

Looking at the per-diet compositional fingerprints (Fig. 2C). We observed an important switch in the community. Indeed, under high fat diet, *Prevotella* were reduced to 3% associated with a dominance in *Streptococcus pasteurianus* (28%) and with a Pseudomonadota bloom. *E.coli* increased from 2% in the control diet to 15% under HF diet. Thus, the proportion of Bacteroidota which represented 33% in animal fed the control diet was reduced to 14%, whilst the proportion of *Bifidobacterium* fell to 5% (instead of 11.2%). It is noteworthy that HF diet gave the lowest evenness of any diet. Twelve and 21 days recovery after 3 days of HF diet resulted in a clearance of the Streptococcus/E. coli bloom, with Bacteroidota/*Prevotella* return to baseline. However, the genus *Bifidobacterium* increased to 18% and Actinomycetota to 22%, associated with increased in *Leuconostoc* and *Megasphaera* (Fig. 2C), thus after 21 days the resilience and return to baseline is not yet achieved. After 3 days of raw meat diet, a enrichment in *Prevotella and* Bacteroidota is observed reaching 31% and 49 % respectively. Interestingly, a rise in *S. equinus* was observed going from 1% to 7.5%, in opposition, dietary *Lactococcus* which were present at 4% in the microbiota of control diet-fed were not detected anymore.

At the functional level (Fig. 3), in sharp contrast to taxonomy, we observed that the predicted functional potential was comparatively stable across diets indicating a signature of functional redundancy. Indeed, alpha and beta diversity of predicted functions show no diet effect in any contrast: no BH-significant alpha contrast both for KEGG and EC, and no overall beta separation (KEGG R² = 0.14, p = 0.79; EC R² = 0.17, p = 0.71; KO R² = 0.19, p = 0.42; all q = 1) (Fig. 3A-C). Differential abundance was sparse and level-dependent. At the pathway (KEGG) level, the most robust functional summary, because pathways aggregate genes, the only significant feature was ko02040 (flagellar assembly), enriched under high-fat (Log2FC +1.90, pcorr 0.049), coherent with the Pseudomonata bloom. At the finer gene levels, a few features reach significance, concentrated in RAW vs CTL (2 EC, 5 KO), mostly low-prevalence genes present in raw but near-absent under control diet, hence inflated fold-changes and in the recovery contrast, CTL post-HF vs HF (1 EC, 5 KO, moderate effects, pcorr ≈ 0.01-0.04). No functional feature reached significance in HF vs CTL at the gene level, nor in COOK vs RAW or recovery vs CTL (Fig. 3D).

Community typing at the function level was dominated by a single signature (∼25 of 31 samples) with only a small high-fat-linked second cluster, far more homogeneous than the two clear taxonomic signatures, so functional composition was markedly more stable across diets than taxonomy, the expected consequence of functional redundancy (Fig. 3E; the EC-gene enterosignature, Fig. S3, shows the same structure). At the KEGG-level, a sPLS-DA VIP ranked discriminant pathways for the five dietary contrasts, but were not validated by nested cross-validation (Fig. S3A); the EC-gene differential- abundance volcano plots were consistent with the sparse gene-level results, i.e. significant EC features only in RAW vs CTL and in recovery vs HF (Fig. S3B); the EC-gene enterosignature reproduces the single-dominant-signature structure (Fig. S3C).

In an attempt to correlate species abundance that were discriminant when comparing the various conditions, we observed several association with physiological conditions including body mass index, glycemia, cholesterol, HDL or triglycerides concentrations (see suppl file for data). Figure S4 illustrates our findings, most of them were not statistically significant when correcting for false discovery rate using Benjamini-Hochberg adjustment. One can notice that some *Parabacteroides* were less abundant in HF diet exposed animal after recovery in control diet were correlated with BMI. Similarly, *Alloprevotella muciniphila* was more abundant when comparing microbiome of the Cook condition *vs* Raw.

## Discussion

In the present paper, we report on the gut microbiota of the grey mouse lemur *M. murinus* from the laboratory breeding population of Brunoy, animal that were born and maintain in captivity among several generation. This small and fast-reproducing primates are interesting model organisms with easy laboratory handling capacities and with a median lifespan of 5.7 years in captivity and a maximum lifespan of 12 years in laboratory-controlled conditions. These animals are interesting species to study metabolism features and aging. They display age-related alterations of their sensory system, motor functions, biological rhythms, immune and endocrine systems, in a similar manner to humans [Languille et al, 2012]. As mouse lemurs age, they are more susceptible to diseases such as neoplasia and sarcopenia [Hämäläinen et al, 2015]. Additionally, their glucoregulatory function changes, similar to what is seen in aged human subjects [Djelti et al, 2016]. Lastly, mouse lemurs exhibit age-related cognitive alterations associated with cerebral atrophy [Picq et al, 2012] as well as amyloid lesions that resemble those observed in Alzheimer’s disease [Mestre-Frances et al, 2000]. Thus, *M. murinus* is an remarkable model to study the role of gut microbiome in physiological and pathophysiological conditions.

To our knowledge the characterization of gut microbiota of grey mouse lemur has never been performed, thus we provide the first gut-microbiome reference for captive grey mouse lemurs, a Bacteroidota/*Prevotella*-dominated community closer to the human than the mouse gut, reinforcing the model’s translational value. Although two studies reported on the gut microbiota of gray-brown mouse lemurs (*M. griseorufus*) that were captured and sampled in Madagascar [Wasimuddin et al, 2019; 2022]. The gut bacterial communities of *M. griseorufus* were dominated by the phyla Bacteroidota (32.3%), Actinomycetota (30.2%), and Bacillota (25.4%), followed by Pseudomonata (5.9%) and Campylobacterota (previously named Epsilonbacteraeota) (4.3%). Other reports on gut microbiota of wild lemurs taxonomically more distant are also available [Greene et al, 2023]. Into the wild, the habitat and food habits are the major discriminant of microbiota composition [Donohue et al, 2022; Wasimuddin et al, 2022]. In the present study, the grey mouse lemur were born and maintained in laboratory breeding conditions and fed a control diet. Their gut microbiota were composed of Bacteroidota (33.0%), Actinomycetota (14.8%), Bacillota (15.6%), Bacillota_C (12.4%), Bacillota_A (11.1%), thus relatively similar to *M. griseorufus*’s microbiota, however Campylobacterota and Pseudomonata were less abundant. Switching from control diet to 3 different diets for only 3 days resulted in microbiota changes showing the flexibility of grey mouse lemur microbiota and reinforcing the interest of this animal as a bridge between rodent model and human physiology that was proposed recently [Camp, 2025]. Indeed, a comparative analysis of our 16S profiles from mouse lemurs showed greater relative similarities with the Unified Human Gastrointestinal Genome (UHGG) catalog [Almeida et al, 2021] than to the mouse reference gut microbiome (MRGM) catalog [Kim et al, 2024] e.g., at the species level: Jaccard 0.035 vs 0.031; precision 0.574 vs 0.124; F1 0.068 vs 0.061. The overlap between *M Microcebus* microbiota and UHGG was 57,4 % and 12,4% toward the MRGM. By contrast, MRGM achieves low absolute coverage of UHGG at the species level (10,7%). Taken together, these findings motivate deeper analyses in *Microcebus* and support its consideration as a pathophysiological model more closely aligned with the human gut microbiome than the mouse. It is noteworthy that UHGG and MRGM are both shotgun metagenomic catalogs, whereas we compared it with 16S rRNA data. A thorough examination of the *M. murinus* gut microbiota composition revealed a *Prevotella* driven enterotype. In human, initially three enterotypes have been revealed, each enterotypes being identifiable by the variation in the abundance of one genus: *Bacteroides* for enterotype 1, *Prevotella* for enterotype 2 and *Ruminococcus* for enterotype 3 [Arumugam et al, 2011]. It has been observed in many studies that the composition of the gut community in healthy adults tends to remain relatively stable over extended periods of time, however, at least for some individuals, gut microbial types may exhibit a degree of fluidity and may not always adhere strictly to clearly defined borders [Costea et al, 2018]. A fourth enterotype, driven by the genus *Bifidobacterium* has been observed in young school-age children, interestingly this enterotype had the lowest gene and species number compared to the others [Zhong et al, 2019]. It is noteworthy that a new analysis of 5,230 human gut metagenomes characterized signatures of bacteria commonly co-occurring, called enterosignatures. Five generalizable enterosignatures were identified, which were predominantly dominated by either *Bacteroides*, Firmicutes (Bacillota), *Prevotella*, *Bifidobacterium*, or *Escherichia* [Frioux et al, 2023]. In the mouse, two enterotypes were described, one dominated by Bacteroidota and *Enterobacteriaceae*, the second one being driven by *Ruminococcus* and *Lachnospiraceae*, interestingly similar to those of the *Bacteroides* and *Ruminococcus* enterotype found in the human population [Hildebrand et al, 2013]. However, in comparison with our findings for which the analyzed animals were from the same facility, the studied mice were from various vendors but maintain in a unique animal facility.

Among the three dietary challenges, high-fat feeding produces the clearest, most interpretable microbiota shift, which reverses on return to the control diet. Indeed, a 3 days HF diet resulted in a reversible *Streptococcus*/Pseudomonata bloom dominated by *S. pasteurianus* (∼28%) and Pseudomonadota (15%) associated with a lower community evenness. Although it was for a short time and isocaloric diet, the microbiota follow the classic direction of high-fat/Western diet dysbiosis and a Proteobacteria expansion observed in human [Hildebrandt et al, 2009; Malesza et al, 2021; Bisanz et al, 2019; Daniel et al, 2014; Shin et al, 2015]. At the functional level, a signal was observed, i.e. enrichment of flagellar assembly (ko02040) which is mechanistically consistent with the Proteobacteria/*Enterobacteriaceae* expansion, tying the taxonomic and predicted-functional observations together. The lower evenness under HF reflected the dominance by a single taxon rather than loss of rare members alone, a common feature of diet-induced dysbiosis. The disturbance caused by a high fat diet was followed by a genuine but imperfect recovery, *S. pasteurianus* and Pseudomonadota returned to baseline, but the community recovered to a level above the baseline rather than reverting to it exactly. Indeed, *Bifidobacterium* and Actinomycetota exceeded control levels with transient *Leuconostoc*/*Megasphaera increased. A* rebound dynamic in which fast- recolonising taxa (e.g. *Bifidobacterium*) transiently overshoot, consistent with recovery being real yet individualized and incomplete [Fassarella et al, 2021; Dethlefsen & Relman, 2011; Leeming et al, 2019], however, the complete individual recovery remained descriptive since only 2 of the 4 HF diet fed animals had full trajectories.

When changing the diet from a control diet to a isocaloric raw beef meat diet and then to a cooked meat diet, we observed changes although the animal were fed this new diet during a short period (3 days). Interestingly, the diet was isocaloric with the same input in macronutrients, only their nature (raw meat or cooked meat) changed. Raw and cooked meat move the community in the opposite direction to high-fat, toward a richer, more *Prevotella*/Bacteroidota-dominated state, with cooking accentuating the raw-diet pattern. The raw meat raised *Prevotella*/Bacteroidota and produced the study’s strongest alpha signal (higher observed richness, p = 0.03-0.05). A meat-based diet enriching *Prevotella* diverges from the canonical animal-based-diet resulting in a *Bacteroides*/bile-tolerant pattern [David et al. 2014] and should be interpreted cautiously: the isocaloric meat preparation is not a pure animal-product diet, and *Prevotella/P. copri* track fibre-fermenting, carbohydrate-rich niches [Wu et al, 2011; Kovatcheva-Datchary et al, 2025; Precup & Vodnar, 2019; De Filippis et al, 2019]. Going to a cooked meat diet accentuated the raw pattern (further *Prevotella* and *P. copri A* enrichment, slightly lower richness). Cooking is an established modifier of microbiome structure and function, so this within-animal comparison is valuable even though no single taxon survived correction [Carmody et al, 2019]. The loss of dietary *Lactococcus* under both meat diets is consistent with removal of a control-diet food microbe rather than a gut-ecological change [Bisanz et al. 2019].

Interestingly, the diet-specific taxonomic turnover was decoupled from an almost invariant predicted-function repertoire signifying a clear signature of functional redundancy. Across every comparison, predicted function barely changed – at the pathway level only flagellar assembly under HD diet, and at the gene level only a handful of mostly low-prevalence features – despite taxonomic turnover, the expected consequence of functional redundancy, whereby many distinct taxa encode the same functions and buffer community function against compositional changes [Louca et al, 2018; Tian et al, 2020; Moya & Ferrer, 2016; Vieira-Silva et al, 2016; Rey-Mariño et al, 2026]. Thus, short-term diet change reshaping taxonomy is well established [David et al, 2014; Turnbaugh et al, 2009]; the novelty here is showing that in this primate the functional layer is buffered even under the HF diet perturbation. However, these functional features are predicted potential (FROGSFUNC/PICRUSt2), not measured; confirmation requires shotgun metagenomics/metatranscriptomics or metabolomics [Douglas et al. 2020]. The co-occurrence networks and keystone candidates are a prioritized hypothesis map, not evidence of interactions. Indeed, Per-signature LUPINE partial-correlation networks, Louvain guilds and Zi-Pi candidate keystones organize the community into interpretable modules, but inferred edges vary with method and threshold and an association is not an interaction [Weiss et al, 2016; Gloor et al, 2017].

The design and analysis are rigorous for the scale, but small paired samples bound what per-feature tests can conclude. Thus, no alpha or single-taxon contrast survived correction and exact sPLS-DA p-values were floored (≥ 0.0625 with 4 pairs). Absence of BH-significant per-feature effects does not demonstrate equivalence, thus the biological signal is carried by community-level composition, typing and the within-animal trajectory. Moreover, sampling spanned the photoperiodic season transition (season retained as a covariate in sensitivity analyses) and only males were studied, so no sex effect was estimable.

## Conclusions

The present study is the first report on the composition of the gut microbiota in grey mouse lemur which are considered a model organism more relevant to human than rodents on a genetical and immunological point of view [Camp, 2025]. Consequently the grey mouse lemur appear to be an excellent model to study metabolic transition and the impact of diet on gut microbiota and the overall animal physiology, and to decipher the mechanisms involved. *M. murinus* harbors a Bacteroidota- /*Prevotella*-dominated gut microbiome aligning more with the human than the mouse reference. Although the study design and the small number of animals is a real caveat. We demonstrated that a short term diet intervention was sufficient to change gut microbiota composition, associated with functional redundancy, indicating that it may also be an pertinent model for microbiota focused studies. The next step will consist in a larger, fully paired replication with complete baseline-challenge- recovery trajectories and independent functional/metabolomic measurements, to convert these exploratory associations into mechanisms.

## Supporting information

Supplementary figures Naour et al

## Acknowledgements

We are most grateful to the Genomics Core Facility GenoA, member of Biogenouest and France Genomique for their technical support. We are grateful to the INRAE MIGALE bioinformatics facility (MIGALE, INRAE, 2020. Migale bioinformatics Facility, doi: 10.15454/1.5572390655343293E12) for computing and storage resources.

## Ethics approval and consent to participate

Experimental procedures have be conducted in accordance with the European Communities Council Directive (86/609/EEC), after their evaluation by an independent ethical council in animal experimentation (CEEA Cuvier n°68) and the agreement by the French Ministry of Higher Education and Research and authorized under the reference #35093-2022020211406097.

## Availability of data and materials

The raw metabarcoding reads generated in this study have been deposited in the NCBI Sequence Read Archive under BioProject accession PRJNAXXXXXX. All individual-specific and sample-specific metadata are available in the Supplementary Table. The full analysis pipeline relies exclusively on publicly available tools and R packages as detailed in the Methods. All custom scripts for data processing, statistical analysis, and figure generation are accessible at: https://github.com/MaximeNaour/

## Authors’ contributions

JT, PP, HMB designed the experiments and supervised the project. FM, JPR, BS, JT did all the animal experiments, IG performed the DNA extraction, MN analyzed and interpreted all data. MN, JT and HMB wrote the manuscript. All authors read and approved the final manuscript.

## Supplementary files

Supplementary figures.

Updated metadata (excel file)

Functional id equivalence (excel file)

## References

Aitchison J. The statistical analysis of compositional data. J R Stat Soc Series B. 1982; 44:139–77. doi: 10.1111/j.2517-6161.1982.tb01195.x.

Almeida A, Nayfach S, Boland M, Strozzi F, Beracochea M, Shi ZJ, et al. A unified catalog of 204,938 reference genomes from the human gut microbiome. Nat Biotechnol. 2021; 39:105–14. doi: 10.1038/s41587-020-0603-3.

Amato KR, Yeoman CJ, Cerda G, Schmitt CA, Cramer JD, Miller ME, Gomez A, Turner TR, Wilson BA, Stumpf RM, Nelson KE, White BA, Knight R, Leigh SR. Variable responses of human and non-human primate gut microbiomes to a Western diet. Microbiome. 2015; 3:53. doi: 10.1186/s40168-015-0120-741.

Anderson MJ. A new method for non-parametric multivariate analysis of variance. Austral Ecol. 2001; 26:32–46. doi: 10.1111/j.1442-9993.2001.01070.pp.x.

Anderson MJ. Distance-based tests for homogeneity of multivariate dispersions. Biometrics. 2006; 62:245–53. doi: 10.1111/j.1541-0420.2005.00440.x.

Andrews S. A Quality Control Tool for High Throughput Sequence Data. 2010. http://www.bioinformatics.babraham.ac.uk/projects/fastqc/.

Arumugam M, Raes J, Pelletier E, Le Paslier D, Yamada T, Mende DR, et al. Enterotypes of the human gut microbiome. Nature. 2011; 473:174–80. doi: 10.1038/nature09944.

Bäckhed F, Ding H, Wang T, Hooper LV, Koh GY, Nagy A, et al. The gut microbiota as an environmental factor that regulates fat storage. Proc Natl Acad Sci U S A. 2004; 101:15718–23. doi: 10.1073/pnas.0407076101.

Bäckhed F, Ley RE, Sonnenburg JL, Peterson DA, Gordon JI. Host-bacterial mutualism in the human intestine. Science. 2005; 307:1915–20. doi: 10.1126/science.1104816.

Bakdash JZ, Marusich LR. Repeated measures correlation. Front Psychol. 2017; 8:456. doi: 10.3389/fpsyg.2017.00456.

Barbera P, Kozlov AM, Czech L, Morel B, Darriba D, Flouri T, et al. EPA-ng: Massively Parallel Evolutionary Placement of Genetic Sequences. Syst Biol. 2019; 68:365–9. doi: 10.1093/sysbio/syy054.

Bates D, Mächler M, Bolker B, Walker S. Fitting linear mixed-effects models using lme4. J Stat Softw. 2015; 67:1–48. doi: 10.18637/jss.v067.i01.

Benjamini Y, Hochberg Y. Controlling the false discovery rate: a practical and powerful approach to multiple testing. J R Stat Soc Series B. 1995; 57:289–300. doi: 10.1111/j.2517-6161.1995.tb02031.x.

Beresford-Jones BS, Forster SC, Stares MD, Notley G, Viciani E, Browne HP, et al. The Mouse Gastrointestinal Bacteria Catalogue enables translation between the mouse and human gut microbiotas via functional mapping. Cell Host Microbe. 2022; 30:124–38.e8. doi: 10.1016/j.chom.2021.12.003.

Bisanz JE, Upadhyay V, Turnbaugh JA, et al. Meta-analysis reveals reproducible gut microbiome alterations in response to a high-fat diet. Cell Host Microbe. 2019;26:265–272.e4. doi:10.1016/j.chom.2019.06.013.

Blondel VD, Guillaume J-L, Lambiotte R, Lefebvre E. Fast unfolding of communities in large networks. J Stat Mech Theory Exp. 2008; 2008:P10008. doi: 10.1088/1742-5468/2008/10/P10008.

Brooks ME, Kristensen K, van Benthem KJ, Magnusson A, Berg CW, Nielsen A, et al. glmmTMB balances speed and flexibility among packages for zero-inflated generalized linear mixed modeling. R J. 2017; 9:378–400. doi: 10.32614/RJ-2017-066.

Callahan BJ, McMurdie PJ, Rosen MJ, Han AW, Johnson AJA, Holmes SP. DADA2: High-resolution sample inference from Illumina amplicon data. Nat Methods. 2016; 13:581–3. doi: 10.1038/nmeth.3869.

Camp JG. The mouse lemur could be science’s next top model. Nature. 2025; 644:43–4. doi: 10.1038/d41586-025-01584-0.

Carmody RN, Bisanz JE, Bowen BP, Maurice CF, Lyalina S, Louie KB, et al. Cooking shapes the structure and function of the gut microbiome. Nat Microbiol. 2019; 4:2052–63. doi: 10.1038/s41564-019-0569-4.

Carter MM, Olm MR, Merrill BD, Dahan D, Tripathi S, Spencer SP, et al. Ultra-deep sequencing of Hadza hunter- gatherers recovers vanishing gut microbes. Cell. 2023; 186:3111–24.e13. doi: 10.1016/j.cell.2023.05.046.

Caspi R, Billington R, Ferrer L, Foerster H, Fulcher CA, Keseler IM, et al. The MetaCyc database of metabolic pathways and enzymes and the BioCyc collection of pathway/genome databases. Nucleic Acids Res. 2016; 44:D471–80. doi: 10.1093/nar/gkv1164.

Caspi R, Billington R, Keseler IM, Kothari A, Krummenacker M, Midford PE, et al. The MetaCyc database of metabolic pathways and enzymes - a 2019 update. Nucleic Acids Res. 2020; 48:D445–53. doi: 10.1093/nar/gkz862.

Costea PI, Hildebrand F, Arumugam M, Bäckhed F, Blaser MJ, Bushman FD, et al. Enterotypes in the landscape of gut microbial community composition. Nat Microbiol. 2018; 3:8–16. doi: 10.1038/s41564-017-0072-8.

Csárdi G, Nepusz T, Traag V, Horvát S, Zanini F, Noom D, et al. igraph: Network Analysis and Visualization. 2006; :2.3.0. doi: 10.32614/CRAN.package.igraph.

Czech L, Barbera P, Stamatakis A. Genesis and Gappa: processing, analyzing and visualizing phylogenetic (placement) data. Bioinformatics. 2020; 36:3263–5. doi: 10.1093/bioinformatics/btaa070.

Dammhahn M, Kappeler PM. Comparative Feeding Ecology of Sympatric *Microcebus berthae* and *M. murinus*. Int J Primatol. 2008; 29:1567–89. doi: 10.1007/s10764-008-9312-3.

Daniel H, Gholami AM, Berry D, et al. High-fat diet alters gut microbiota physiology in mice. ISME J. 2014;8:295–308. doi:10.1038/ismej.2013.155

David LA, Maurice CF, Carmody RN, et al. Diet rapidly and reproducibly alters the human gut microbiome. Nature. 2014;505:559–563. doi:10.1038/nature12820

De Filippis F, Pasolli E, Tett A, et al. Distinct genetic and functional traits of human intestinal Prevotella copri strains are associated with different habitual diets. Cell Host Microbe. 2019;25:444–453.e3. doi:10.1016/j.chom.2019.01.004

Dethlefsen L, Relman DA. Incomplete recovery and individualized responses of the human distal gut microbiota to repeated antibiotic perturbation. Proc Natl Acad Sci USA. 2011;108(Suppl 1):4554–4561. doi:10.1073/pnas.1000087107

Djelti F, Dhenain M, Terrien J, Picq JL, Hardy I, Champeval D, et al. Impaired fasting blood glucose is associated to cognitive impairment and cerebral atrophy in middle-aged non-human primates. Aging (Albany NY). 2016; 9:173–86. doi: 10.18632/aging.101148.

Donohue ME, Rowe AK, Kowalewski E, et al. Significant effects of host dietary guild and phylogeny in wild lemur gut microbiomes. ISME Commun. 2022;2:33. doi:10.1038/s43705-022-00115-6

Doré J, Blottière HM. The influence of diet on the gut microbiota and its consequences for health. Curr Opin Biotechnol. 2015; 32: 195–9. doi: 10.1016/j.copbio.2015.01.002.

Douglas GM, Maffei VJ, Zaneveld JR, Yurgel SN, Brown JR, Taylor CM, et al. PICRUSt2 for prediction of metagenome functions. Nat Biotechnol. 2020; 38:685–8. doi: 10.1038/s41587-020-0548-6.

Eddy SR. Accelerated Profile HMM Searches. PLoS Comput Biol. 2011; 7:e1002195. doi: 10.1371/journal.pcbi.1002195.

Edgar RC, Haas BJ, Clemente JC, Quince C, Knight R. UCHIME improves sensitivity and speed of chimera detection. Bioinformatics. 2011; 27:2194–200. doi: 10.1093/bioinformatics/btr381.

Escudié F, Auer L, Bernard M, Mariadassou M, Cauquil L, Vidal K, et al. FROGS: Find, Rapidly, OTUs with Galaxy Solution. Bioinformatics. 2018; 34:1287–94. Doi: 10.1093/bioinformatics/btx791.

Ewels P, Magnusson M, Lundin S, Käller M. MultiQC: summarize analysis results for multiple tools and samples in a single report. Bioinformatics. 2016; 32:3047–8. doi:10.1093/bioinformatics/btw354.

Ezran C, Liu S, Chang S, Ming J, Guethlein LA, Wang MFZ, et al. Mouse lemur cell atlas informs primate genes, physiology and disease. Nature. 2025; 644: 185–96. doi: 10.1038/s41586-025-09114-8.

Fassarella M, Blaak EE, Penders J, et al. Gut microbiome stability and resilience: elucidating the response to perturbations in order to modulate gut health. Gut. 2021;70:595–605. doi:10.1136/gutjnl-2020-321747.

Ferrari SLP, Cribari-Neto F. Beta regression for modelling rates and proportions. J Appl Stat. 2004; 31:799–815. doi: 10.1080/0266476042000214501.

Frioux C, Ansorge R, Özkurt E, Ghassemi Nedjad C, Fritscher J, Quince C, et al. Enterosignatures define common bacterial guilds in the human gut microbiome. Cell Host Microbe. 2023; 31:1111–25.e6. doi: 10.1016/j.chom.2023.05.024.

Gloor GB, Macklaim JM, Pawlowsky-Glahn V, Egozcue JJ. Microbiome datasets are compositional: and this is not optional. Front Microbiol. 2017;8:2224. doi:10.3389/fmicb.2017.02224

Greene LK, McKenney EA, Gasper W, Wrampelmeier C, Hayer S, Ehmke EE, Clayton JB. Gut Site and Gut Morphology Predict Microbiome Structure and Function in Ecologically Diverse Lemurs. Microb Ecol. 2023; 85:1608–19. doi: 10.1007/s00248-022-02034-4.

Gu Z, Eils R, Schlesner M. Complex heatmaps reveal patterns and correlations in multidimensional genomic data. Bioinformatics. 2016; 32:2847–9. doi: 10.1093/bioinformatics/btw313.

Guimerà R, Amaral LAN. Functional cartography of complex metabolic networks. Nature. 2005; 433:895–900. doi: 10.1038/nature03288.

Hämäläinen A, Dammhahn M, Aujard F, Kraus C. Losing grip: Senescent decline in physical strength in a small- bodied primate in captivity and in the wild. Exp Gerontol. 2015; 61:54–61. doi: 10.1016/j.exger.2014.11.017.

Hauffe HC, Barelli C. Conserve the germs: the gut microbiota and adaptive potential. Conserv Genet. 2019; 20:19–27.doi: 10.1007/s10592-019-01150-y.

Hildebrand F, Nguyen TL, Brinkman B, Yunta RG, Cauwe B, Vandenabeele P, et al. Inflammation-associated enterotypes, host genotype, cage and inter-individual effects drive gut microbiota variation in common laboratory mice. Genome Biol. 2013; 14:R4. doi: 10.1186/gb-2013-14-1-r4.

Hildebrandt MA, Hoffmann C, Sherrill-Mix SA, et al. High-fat diet determines the composition of the murine gut microbiome independently of obesity. Gastroenterology. 2009;137:1716–1724.e2. doi:10.1053/j.gastro.2009.08.042

Kaczmarek JL, Musaad SM, Holscher HD. Time of day and eating behaviors are associated with the composition and function of the human gastrointestinal microbiota. Am J Clin Nutr. 2017;106:1220–31. doi: 10.3945/ajcn.117.156380.

Kanehisa M, Furumichi M, Sato Y, Matsuura Y, Ishiguro-Watanabe M. KEGG: biological systems database as a model of the real world. Nucleic Acids Res. 2025; 53:D672–7. doi: 10.1093/nar/gkae909.

Kim N, Kim CY, Ma J, Yang S, Park DJ, Ha SJ, Belenky P, Lee I. MRGM: an enhanced catalog of mouse gut microbial genomes substantially broadening taxonomic and functional landscapes. Gut Microbes. 2024; 16:2393791. doi: 10.1080/19490976.2024.2393791.

Kodikara S, Lê Cao K-A. Microbial network inference for longitudinal microbiome studies with LUPINE. Microbiome. 2025; 13:64. doi: 10.1186/s40168-025-02041-w.

Kovatcheva-Datchary P, Nilsson A, Akrami R, et al. Dietary fiber-induced improvement in glucose metabolism is associated with increased abundance of Prevotella. Cell Metab. 2015;22:971–982. doi:10.1016/j.cmet.2015.10.001.

Languille S, Blanc S, Blin O, Canale CI, Dal-Pan A, Devau G, et al. The grey mouse lemur: a non-human primate model for ageing studies. Ageing Res Rev. 2012; 11:150–62. doi: 10.1016/j.arr.2011.07.001.

Lê Cao K-A, Boitard S, Besse P. Sparse PLS discriminant analysis: biologically relevant feature selection and graphical displays for multiclass problems. BMC Bioinformatics. 2011; 12:253. doi: 10.1186/1471-2105-12-253.

Lee DD, Seung HS. Learning the parts of objects by non-negative matrix factorization. Nature. 1999; 401:788–91. doi: 10.1038/44565.

Leeming ER, Johnson AJ, Spector TD, Le Roy CI. Effect of diet on the gut microbiota: rethinking intervention duration. Nutrients. 2019;11:2862. doi:10.3390/nu11122862.

Lenth RV. emmeans: Estimated Marginal Means, aka Least-Squares Means. 2025; :2.0.2. doi: 10.32614/CRAN.package.emmeans.

Louca S, Doebeli M. Efficient comparative phylogenetics on large trees. Bioinformatics. 2018; 34:1053–5. doi: 10.1093/bioinformatics/btx701.

Louca S, Polz MF, Mazel F, et al. Function and functional redundancy in microbial systems. Nat Ecol Evol. 2018;2:936–943. doi:10.1038/s41559-018-0519-1.

Malesza IJ, Malesza M, Walkowiak J, et al. High-fat, Western-style diet, systemic inflammation, and gut microbiota: a narrative review. Cells. 2021;10:3164. doi:10.3390/cells10113164.

Martino C, Morton JT, Marotz CA, Thompson LR, Tripathi A, Knight R, et al. A novel sparse compositional technique reveals microbial perturbations. mSystems. 2019; 4:e00016–19. doi: 10.1128/mSystems.00016-19.

McMurdie PJ, Holmes S. phyloseq: An R Package for Reproducible Interactive Analysis and Graphics of Microbiome Census Data. PLoS one. 2013; 8:e61217. doi: 10.1371/journal.pone.0061217.

Mestre-Francés N, Keller E, Calenda A, Barelli H, Checler F, Bons N. Immunohistochemical analysis of cerebral cortical and vascular lesions in the primate Microcebus murinus reveal distinct amyloid beta1-42 and beta1-40 immunoreactivity profiles. Neurobiol Dis. 2000; 7:1–8. doi: 10.1006/nbdi.1999.0270.

Monti S, Tamayo P, Mesirov J, Golub T. Consensus clustering: a resampling-based method for class discovery and visualization of gene expression microarray data. Mach Learn. 2003; 52:91–118. doi: 10.1023/A:1023949509487.

Moon KR, van Dijk D, Wang Z, Gigante S, Burkhardt DB, Chen WS, et al. Visualizing structure and transitions in high-dimensional biological data. Nat Biotechnol. 2019; 37:1482–92. doi: 10.1038/s41587-019-0336-3.

Moya A, Ferrer M. Functional redundancy-induced stability of gut microbiota subjected to disturbance. Trends Microbiol. 2016;24:402–413. doi:10.1016/j.tim.2016.02.002

Nguyen TL, Vieira-Silva S, Liston A, Raes J. How informative is the mouse for human gut microbiota research? Dis Model Mech. 2015; 8:1–16. doi: 10.1242/dmm.017400.

O’Keefe SJ. The association between dietary fibre deficiency and high-income lifestyle-associated diseases: Burkitt’s hypothesis revisited. Lancet Gastroenterol Hepatol. 2019; 4:984–96. doi: 10.1016/S2468-1253(19)30257-2.

Oksanen J, Simpson GL, Blanchet FG, Kindt R, Legendre P, Minchin PR, et al. vegan: Community Ecology Package. 2025; :2.7–1. doi: 10.32614/CRAN.package.vegan.

Owen AB, Perry PO. Bi-cross-validation of the SVD and the nonnegative matrix factorization. Ann Appl Stat. 2009; 3:564–94. doi: 10.1214/08-AOAS227.

Palarea-Albaladejo J, Martín-Fernández JA. zCompositions - R package for multivariate imputation of left- censored data under a compositional approach. Chemometr Intell Lab Syst. 2015; 143:85–96. doi: 10.1016/j.chemolab.2015.02.019.

Pedersen TL. ggraph: An Implementation of Grammar of Graphics for Graphs and Networks. 2024; :2.2.1. doi: 10.32614/CRAN.package.ggraph.

Perret, M., Aujard, F. Regulation by Photoperiod of Seasonal Changes in Body Mass and Reproductive Function in Gray Mouse Lemurs (*Microcebus murinus*): Differential Responses by Sex. Int J Primatol. 2001; 22: 5–24.doi:10.1023/A:1026457813626.

Picq JL, Aujard F, Volk A, Dhenain M. Age-related cerebral atrophy in nonhuman primates predicts cognitive impairments. Neurobiol Aging. 2012; 33:1096–109. doi: 10.1016/j.neurobiolaging.2010.09.009.

Pielou EC. The measurement of diversity in different types of biological collections. J Theor Biol. 1966; 13:131–44. doi: 10.1016/0022-5193(66)90013-0.

Precup G, Vodnar DC. Gut Prevotella as a possible biomarker of diet and its eubiotic versus dysbiotic roles. Br J Nutr. 2019;122:131–140. doi:10.1017/S0007114519000680

Preguiça I, Alves A, Nunes S, Fernandes R, Gomes P, Viana SD, Reis F. Diet-induced rodent models of obesity- related metabolic disorders-A guide to a translational perspective. Obes Rev. 2020; 21:e13081. doi: 10.1111/obr.13081.

R Core Team. R: A Language and Environment for Statistical Computing. 2024. https://www.R-project.org/.

Radespiel U, Reimann W, Rahelinirina M, Zimmermann E. Feeding Ecology of Sympatric Mouse Lemur Species in Northwestern Madagascar. Int J Primatol. 2006; 27:311–21. doi: 10.1007/s10764-005-9005-0.

Rey-Mariño A, et al. Patterns of gut microbiome composition, function and dynamics across age groups over a three-year period (functional stability exceeds taxonomic stability). Front Microbiol. 2026. (in press)

Riedl RA, Atkinson SN, Burnett CML, Grobe JL, Kirby JR. The Gut Microbiome, Energy Homeostasis, and Implications for Hypertension. Curr Hypertens Rep. 2017; 19:27. doi: 10.1007/s11906-017-0721-6.

Rina Evasoa M, Radespiel U, Hasiniaina AF, Rasoloharijaona S, Randrianambinina B, Rakotondravony R, Zimmermann E. Variation in reproduction of the smallest-bodied primate radiation, the mouse lemurs (*Microcebus spp.*): A synopsis. Am J Primatol. 2018; 80:e22874. doi: 10.1002/ajp.22874.

Rohart F, Gautier B, Singh A, Cao K-AL. mixOmics: An R package for ‘omics feature selection and multiple data integration. PLoS Comput Biol. 2017; 13:e1005752. doi: 10.1371/journal.pcbi.1005752.

Rousseeuw PJ. Silhouettes: a graphical aid to the interpretation and validation of cluster analysis. J Comput Appl Math. 1987; 20:53–65. doi: 10.1016/0377-0427(87)90125-7.

Scheithauer TPM, Dallinga-Thie GM, de Vos WM, Nieuwdorp M, van Raalte DH. Causality of small and large intestinal microbiota in weight regulation and insulin resistance. Mol Metab. 2016; 5:759–70. doi: 10.1016/j.molmet.2016.06.002.

Schmid J, Speakman JR. Daily energy expenditure of the grey mouse lemur (*Microcebus murinus*): a small primate that uses torpor. J Comp Physiol B. 2000; 170:633–41. doi: 10.1007/s003600000146.

Shin NR, Whon TW, Bae JW. Proteobacteria: microbial signature of dysbiosis in gut microbiota. Trends Biotechnol. 2015;33:496–503. doi:10.1016/j.tibtech.2015.06.011.

Spearman C. The proof and measurement of association between two things. Am J Psychol. 1904; 15:72–101. doi: 10.2307/1412159.

Tabula Microcebus Consortium; Ezran C, Liu S, Chang S, Ming J, Botvinnik O, et al. A molecular cell atlas of mouse lemur, an emerging model primate. Nature. 2025; 644, pages 173–184. doi: 10.1038/s41586-025-09113-9.

Tian L, Wang X-W, Wu A-K, et al. Deciphering functional redundancy in the human microbiome. Nat Commun. 2020;11:6217. doi:10.1038/s41467-020-19940-1

Turnbaugh PJ, Ley RE, Mahowald MA, Magrini V, Mardis ER, Gordon JI. An obesity-associated gut microbiome with increased capacity for energy harvest. Nature. 2006; 444:1027–31. doi: 10.1038/nature05414.

Turnbaugh PJ, Ridaura VK, Faith JJ, et al. The effect of diet on the human gut microbiome: a metagenomic analysis in humanized gnotobiotic mice. Sci Transl Med. 2009;1:6ra14. doi:10.1126/scitranslmed.3000322

Vieira-Silva S, Falony G, Darzi Y, et al. Species-function relationships shape ecological properties of the human gut microbiome. Nat Microbiol. 2016;1:16088. doi:10.1038/nmicrobiol.2016.88

Wang Q, Garrity GM, Tiedje JM, Cole JR. Naïve Bayesian Classifier for Rapid Assignment of rRNA Sequences into the New Bacterial Taxonomy. Appl Environ Microbiol. 2007; 73:5261–7. doi: 10.1128/AEM.00062-07.

Wang Y, Kuang Z, Yu X, Kelly A. Ruhn KA, Kubo M, Hooper LV. The intestinal microbiota regulates body composition through NFIL3 and the circadian clock. Science. 2017; 357:912–6. doi: 10.1126/science.aan067.

Wasimuddin, Corman VM, Ganzhorn JU, Rakotondranary J, Ratovonamana YR, Drosten C, Sommer S. Adenovirus infection is associated with altered gut microbial communities in a non-human primate. Sci Rep. 2019; 9:13410. doi: 10.1038/s41598-019-49829-z.

Wasimuddin, Malik H, Ratovonamana YR, Rakotondranary SJ, Ganzhorn JU, Sommer S. Anthropogenic Disturbance Impacts Gut Microbiome Homeostasis in a Malagasy Primate. Front Microbiol. 2022; 13:911275. doi: 10.3389/fmicb.2022.911275.

Weiss S, Van Treuren W, Lozupone C, et al. Correlation detection strategies in microbial data sets vary widely in sensitivity and precision. ISME J. 2016;10:1669-1681. doi:10.1038/ismej.2015.235

Wright PC. Lemur traits and Madagascar ecology: coping with an island environment. Am J Phys Anthropol. 1999; 29:31–72. doi: 10.1002/(sici)1096-8644(1999)110:29+<31::aid-ajpa3>3.0.co;2-0.

Wu GD, Chen J, Hoffmann C, et al. Linking long-term dietary patterns with gut microbial enterotypes. Science. 2011;334:105–108. doi:10.1126/science.1208344

Xiao L, Feng Q, Liang S, Sonne SB, Xia Z, Qiu X, et al. A catalog of the mouse gut metagenome. Nat Biotechnol. 2015; 33:1103–8. doi: 10.1038/nbt.3353.

Ye Y, Doak TG. A Parsimony Approach to Biological Pathway Reconstruction/Inference for Genomes and Metagenomes. PLOS Comput Biol. 2009; 5:e1000465. doi: 10.1371/journal.pcbi.1000465.

Zhang J, Kobert K, Flouri T, Stamatakis A. PEAR: a fast and accurate Illumina Paired-End reAd mergeR. Bioinformatics. 2014; 30:614–20. Doi: 10.1093/bioinformatics/btt593.

Zhong H, Penders J, Shi Z, Ren H, Cai K, Fang C, et al. Impact of early events and lifestyle on the gut microbiota and metabolic phenotypes in young school-age children. Microbiome. 2019; 7:2. doi: 10.1186/s40168-018-0608-z.

Zhou H, He K, Chen J, Zhang X. LinDA: linear models for differential abundance analysis of microbiome compositional data. Genome Biol. 2022; 23:95. doi: 10.1186/s13059-022-02655-5.

