## Supplementary figures Naour et al for "Grey mouse lemurs, *Microcebus murinus,* are a relevant model to study gut microbiome flexibility in response to diet changes"

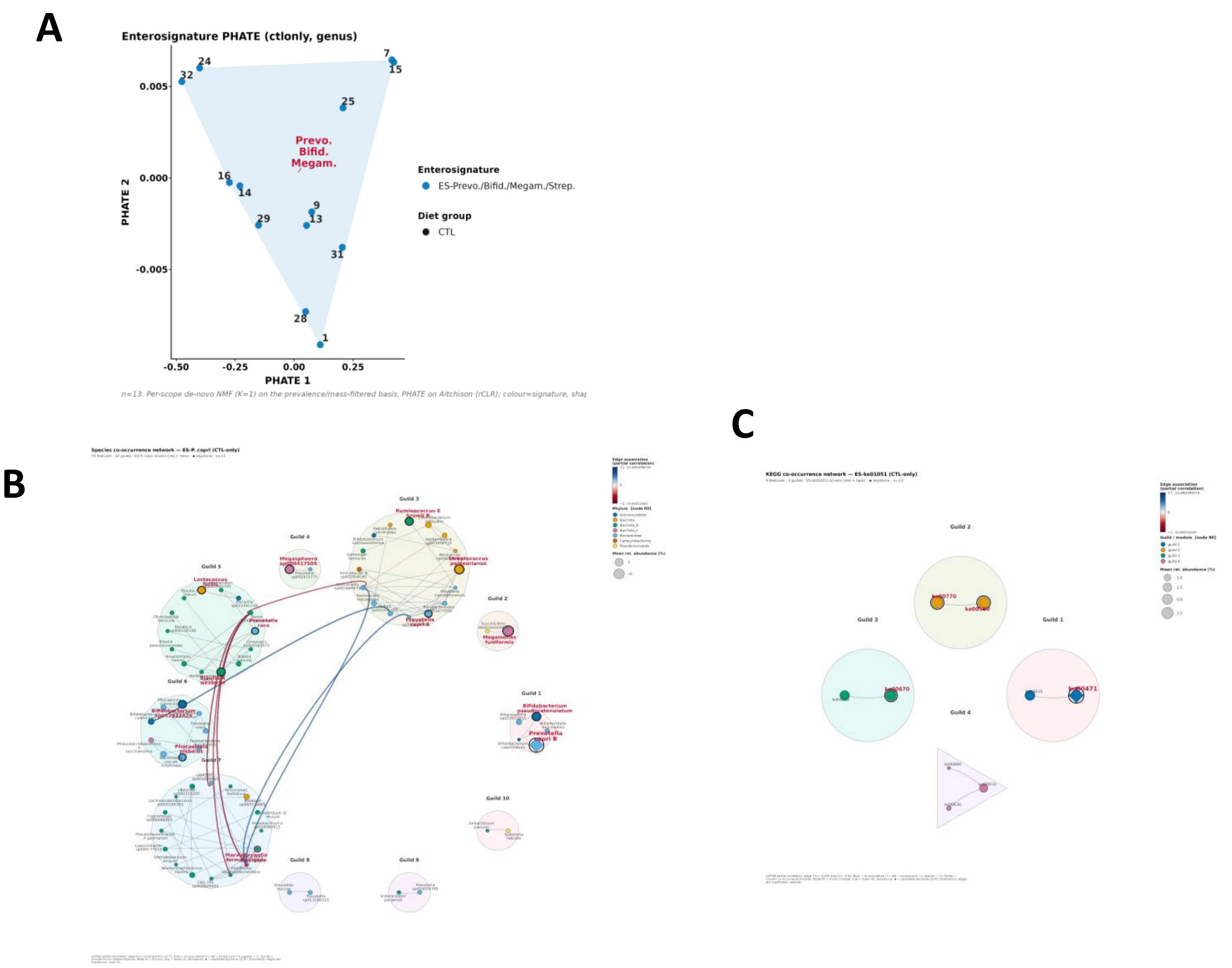


**Figure S1 (Fig. 1). Community structure of the control-diet microbiome: genus enterosignature and its co-occurrence networks.** A) Enterosignature structure at the genus level (control animals only, PHATE embedding); B) Co-occurrence network of the principal species-level signature (ES-*P. copri*): nodes are species coloured by phylum and sized by mean relative abundance; edges are LUPINE partial correlations (blue = co-abundance, red = co-exclusion); shaded hulls are Louvain guilds; a diamond marks a candidate keystone; signature driver species are labelled in red; C) Co-occurrence network of the principal KEGG-pathway signature (ES-ko01051), same conventions with pathways as nodes.

**Notes.** Co-occurrence network: LUPINE partial-correlation graph in which an edge is a conditional association between two features (independent of the others); edge opacity scales with strength.

Guild: a Louvain co-occurrence module (a group of features that co-vary together).

Candidate keystone: a Guimera-Amaral Zi-Pi hub or connector node; exploratory, not a validated keystone.

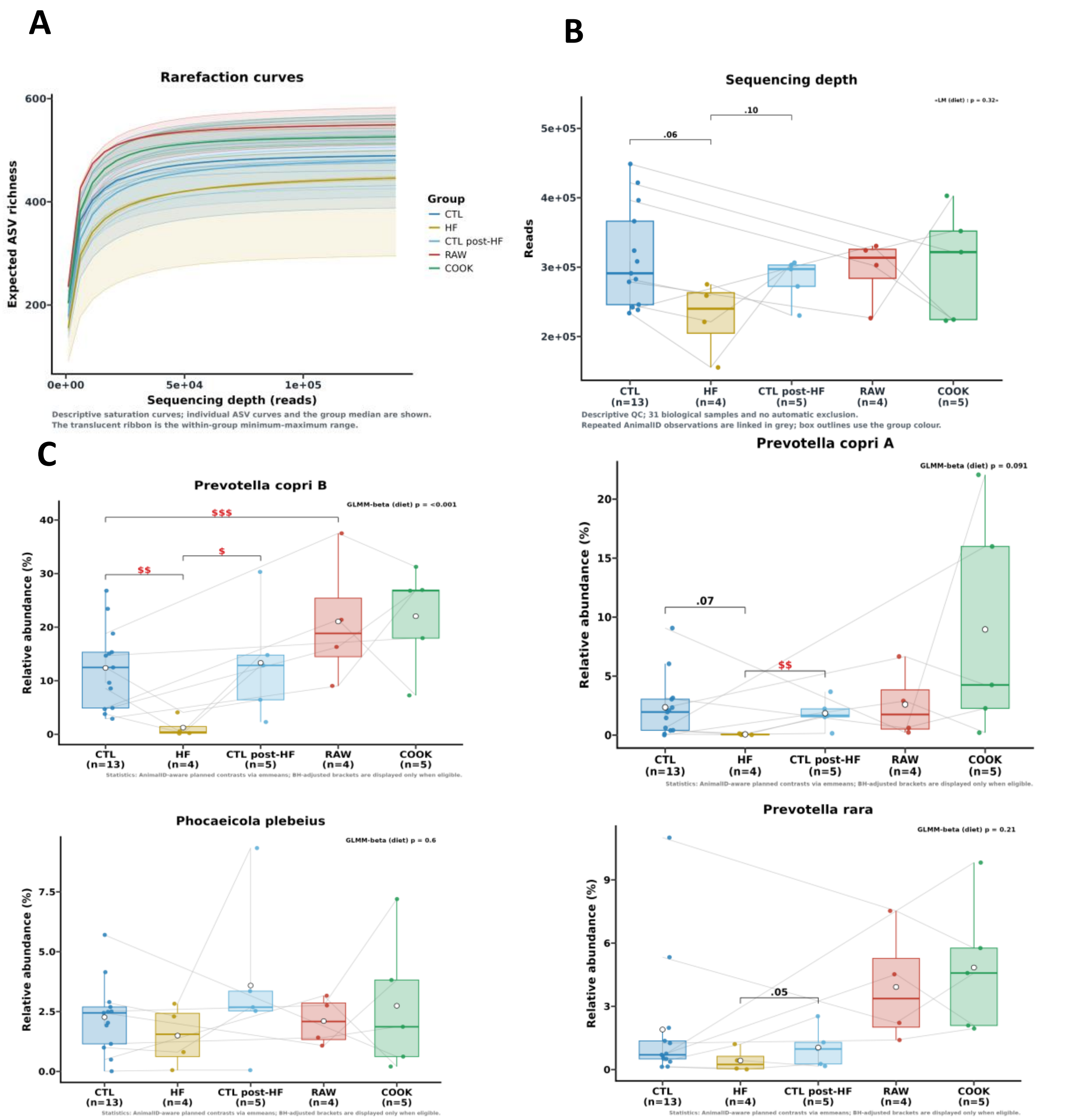


**Figure S2. Sequencing quality and top-taxa dynamics supporting Figure 2**. A) Rarefaction curves per diet group (richness versus sequencing effort), confirming that sampling depth captured the community; B) Sequencing-depth boxplots per diet group; C) Relative abundance of the top 4 most abundant species across all samples : *Prevotella* copri B, *P. copri A*, *Phocaeicola plebeius* and *P. rara*;

**Figure S2 D (below). Supervised discrimination of diets at the species level.** Sparse PLS-DA VIP-importance bar plots for the five diet contrasts (HF vs CTL, CTL post-HF vs HF, CTL post-HF vs CTL, RAW vs CTL, COOK vs RAW); bars rank the species with the highest Variable Importance in Projection, i.e. those contributing most to separating the two diets in a descriptive full-data sPLS-DA.

**Notes.** sPLS-DA VIP: Variable Importance in Projection from a sparse partial-least-squares discriminant analysis; higher VIP means the feature contributes more to separating the two diets.

D

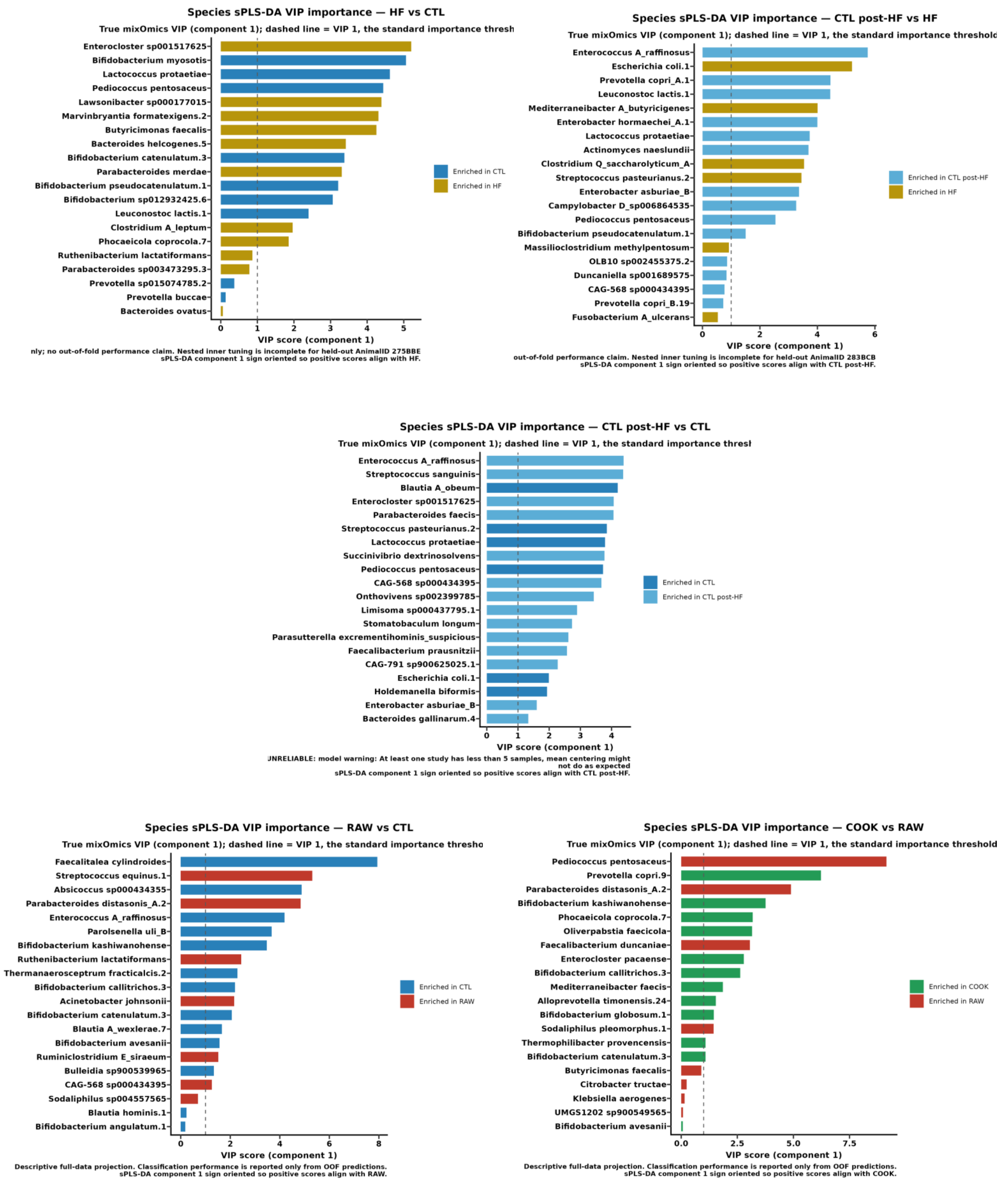

Species sPLS-DA VIP importance — HF vs CTL
True mixOmics VIP (component 1); dashed line = VIP 1, the standard importance thresh
Enterocloster sp001517625
Bifidobacterium myosotis
Lactococcus protaetiae
Pediococcus pentosaceus
Lawsonibacter sp000177015
Marvinbryantia formatexigens.2
Butyricimonas faecalis
Bacteroides helcogenes.5
Bifidobacterium catenulatum.3
Parabacteroides merdae
Bifidobacterium pseudocatenulatum.1
Bifidobacterium sp012932425.6
Leuconostoc lactis.1
Clostridium A_leptum
Phocaeicola coprocola.7
Ruthenibacterium lactatiformans
Parabacteroides sp003473295.3
Prevotella sp015074785.2
Prevotella buccae
Bacteroides ovatus
0 1 2 3 4 5
VIP score (component 1)
Enriched in CTL
Enriched in HF
nly; no out-of-fold performance claim. Nested inner tuning is incomplete for held-out AnimalID 275BBE
sPLS-DA component 1 sign oriented so positive scores align with HF.
Species sPLS-DA VIP importance — CTL post-HF vs HF
True mixOmics VIP (component 1); dashed line = VIP 1, the standard importance threshold
Enterococcus A_raffinosus
Escherichia coli.1
Prevotella copri_A.1
Leuconostoc lactis.1
Mediterraneibacter A_butyricigenes
Enterobacter hormaechei_A.1
Lactococcus protaetiae
Actinomyces naeslundii
Clostridium Q_saccharolyticum_A
Streptococcus pasteurianus.2
Enterobacter asburiae_B
Campylobacter D_sp006864535
Pediococcus pentosaceus
Bifidobacterium pseudocatenulatum.1
Massilioclostridium methylpentosum
OLB10 sp002455375.2
Duncaniella sp001689575
CAG-568 sp000434395
Prevotella copri_B.19
Fusobacterium A_ulcerans
0 2 4 6
VIP score (component 1)
Enriched in CTL post-HF
Enriched in HF
out-of-fold performance claim. Nested inner tuning is incomplete for held-out AnimalID 283BCB
sPLS-DA component 1 sign oriented so positive scores align with CTL post-HF.
Species sPLS-DA VIP importance — CTL post-HF vs CTL
True mixOmics VIP (component 1); dashed line = VIP 1, the standard importance thresl
Enterococcus A_raffinosus
Streptococcus sanguinis
Blautia A_obeum
Enterocloster sp001517625
Parabacteroides faecis
Streptococcus pasteurianus.2
Lactococcus protaetiae
Succinivibrio dextrinosolvens
Pediococcus pentosaceus
CAG-568 sp000434395
Onthovivens sp002399785
Limisoma sp000437795.1
Stomatobaculum longum
Parasutterella excrementihominis_suspicious
Faecalibacterium prausnitzii
CAG-791 sp900625025.1
Escherichia coli.1
Holdemanella biformis
Enterobacter asburiae_B
Bacteroides gallinarum.4
0 1 2 3 4
VIP score (component 1)
Enriched in CTL
Enriched in CTL post-HF
JNRELIABLE: model warning: At least one study has less than 5 samples, mean centering might
not do as expected
sPLS-DA component 1 sign oriented so positive scores align with CTL post-HF.
Species sPLS-DA VIP importance — RAW vs CTL
True mixOmics VIP (component 1); dashed line = VIP 1, the standard importance thresho
Faecalitalea cylindroides
Streptococcus equinus.1
Absicoccus sp000434355
Parabacteroides distasonis_A.2
Enterococcus A_raffinosus
Parolsenella uli_B
Bifidobacterium kashiwanohense
Ruthenibacterium lactatiformans
Thermanaerosceptrum fracticalcis.2
Bifidobacterium callitrichos.3
Acinetobacter johnsonii
Bifidobacterium catenulatum.3
Blautia A_wexlerae.7
Bifidobacterium avesanii
Ruminiclostridium E_siraeum
Bulleidia sp900539965
CAG-568 sp000434395
Sodaliphilus sp004557565
Blautia hominis.1
Bifidobacterium angulatum.1
0 2 4 6 8
VIP score (component 1)
Enriched in CTL
Enriched in RAW
Descriptive full-data projection. Classification performance is reported only from OOF predictions.
sPLS-DA component 1 sign oriented so positive scores align with RAW.
Species sPLS-DA VIP importance — COOK vs RAW
True mixOmics VIP (component 1); dashed line = VIP 1, the standard importance threshold
Pediococcus pentosaceus
Prevotella copri.9
Parabacteroides distasonis_A.2
Bifidobacterium kashiwanohense
Phocaeicola coprocola.7
Oliverpabstia faecicola
Faecalibacterium duncaniae
Enterocloster pacaense
Bifidobacterium callitrichos.3
Mediterraneibacter faecis
Alloprevotella timonensis.24
Bifidobacterium globosum.1
Sodaliphilus pleomorphus.1
Thermophilibacter provencensis
Bifidobacterium catenulatum.3
Butyricimonas faecalis
Citrobacter tructae
Klebsiella aerogenes
UMGS1202 sp900549565
Bifidobacterium avesanii
0.0 2.5 5.0 7.5
VIP score (component 1)
Enriched in COOK
Enriched in RAW
Descriptive full-data projection. Classification performance is reported only from OOF predictions.
sPLS-DA component 1 sign oriented so positive scores align with COOK.

A

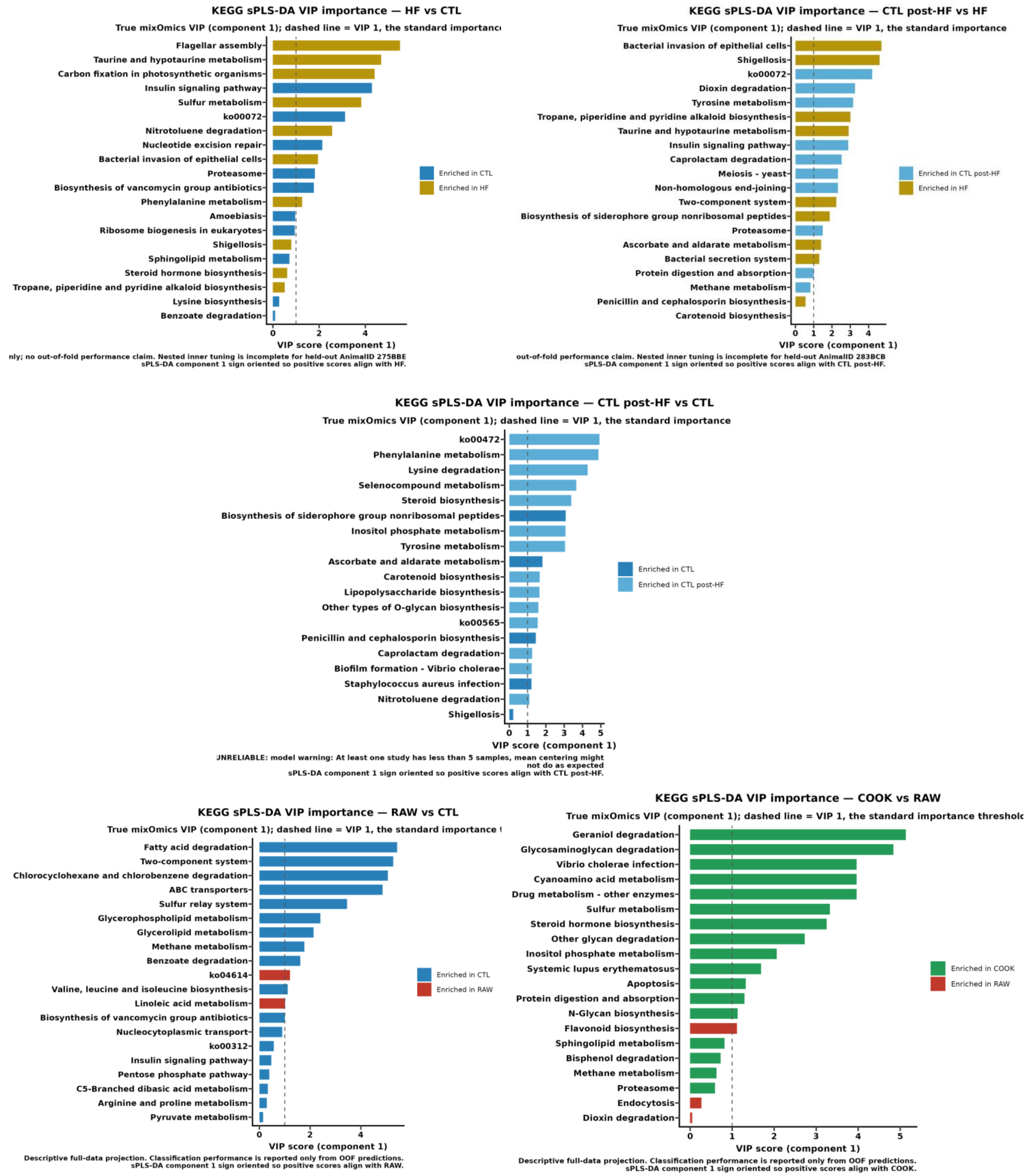


**Figure S3. A Supervised discrimination of diets at the KEGG-pathway level.** Sparse PLS-DA VIP-importance bar plots for the five diet contrasts, ranking the pathways that contribute most to separating each diet pair (descriptive full-data sPLS-DA).

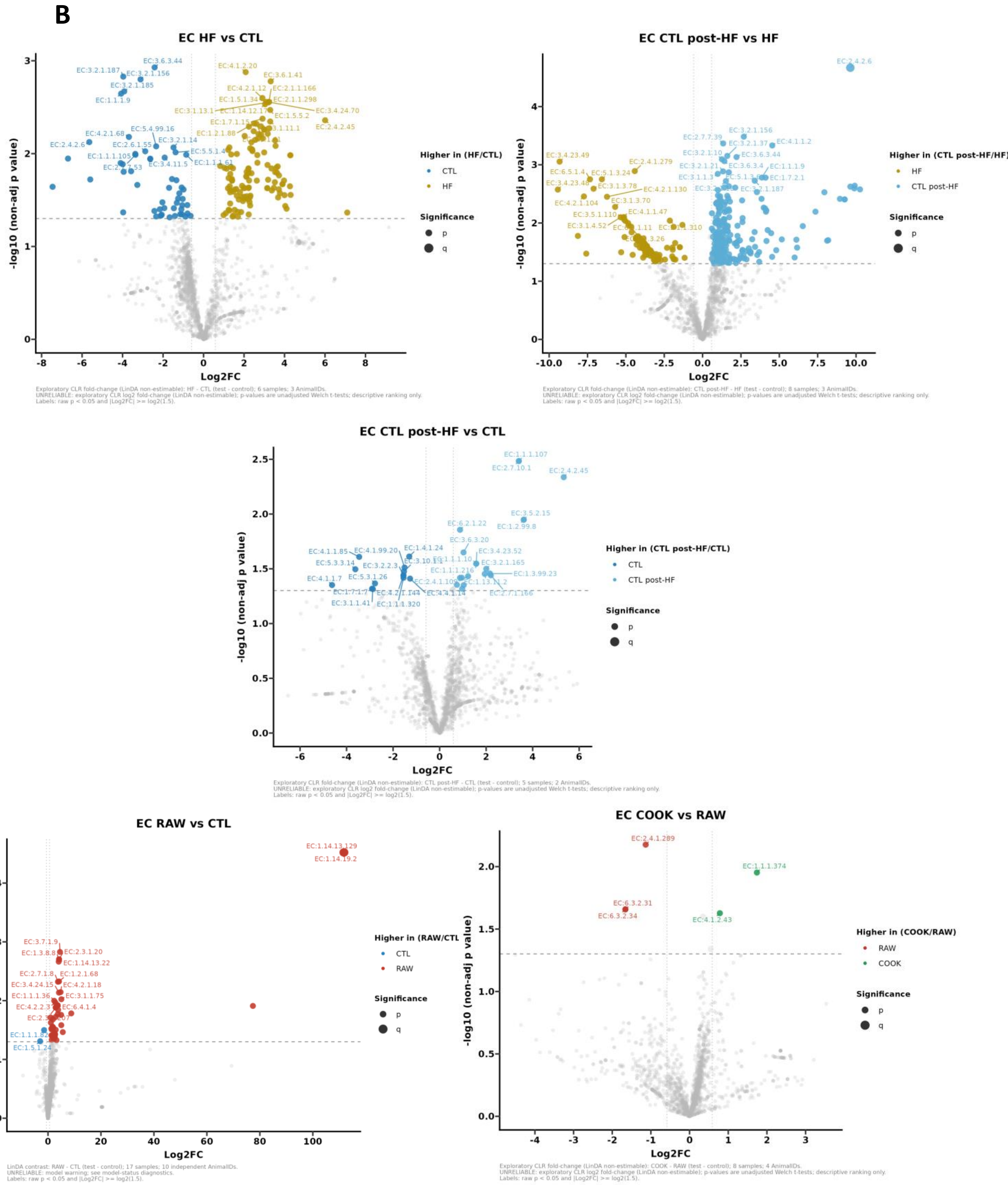


**Figure S3 B). EC-gene differential abundance across diets.** LinDA differential abundance at the EC-gene level for the five diet contrasts, shown as volcano plots (log2 fold-change of the test diet versus its reference on the x-axis against -log10 p on the y-axis; features passing the significance threshold are highlighted).

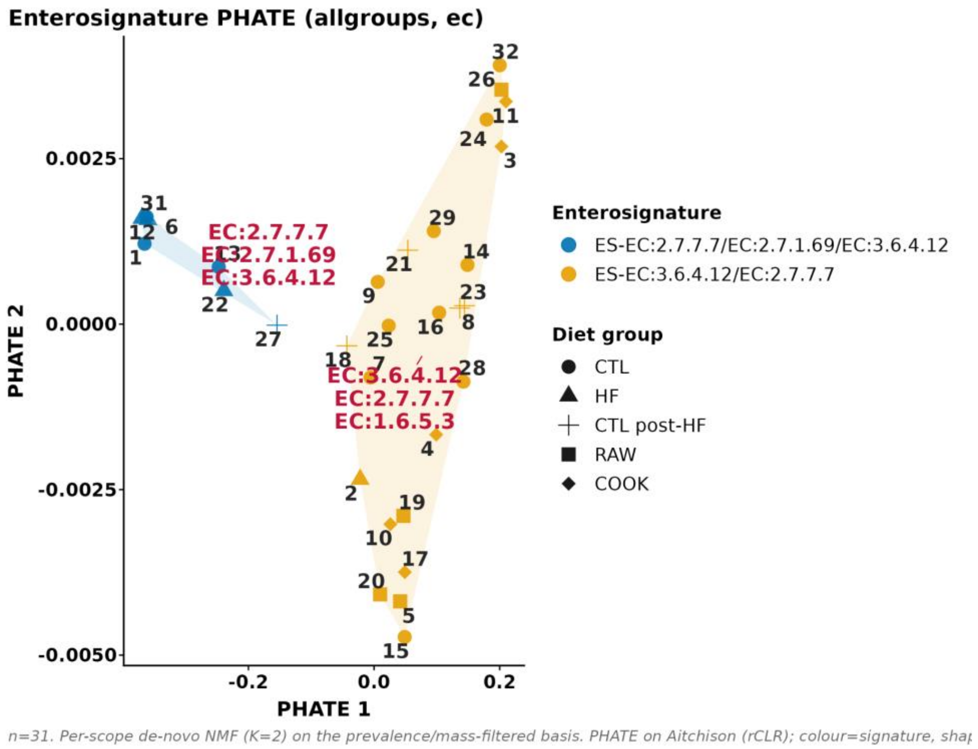


**Figure S3 C. EC-gene enterosignature.** Enterosignature structure at the EC-gene level across all diets (PHATE embedding); points are samples coloured by signature and shaped by diet group.

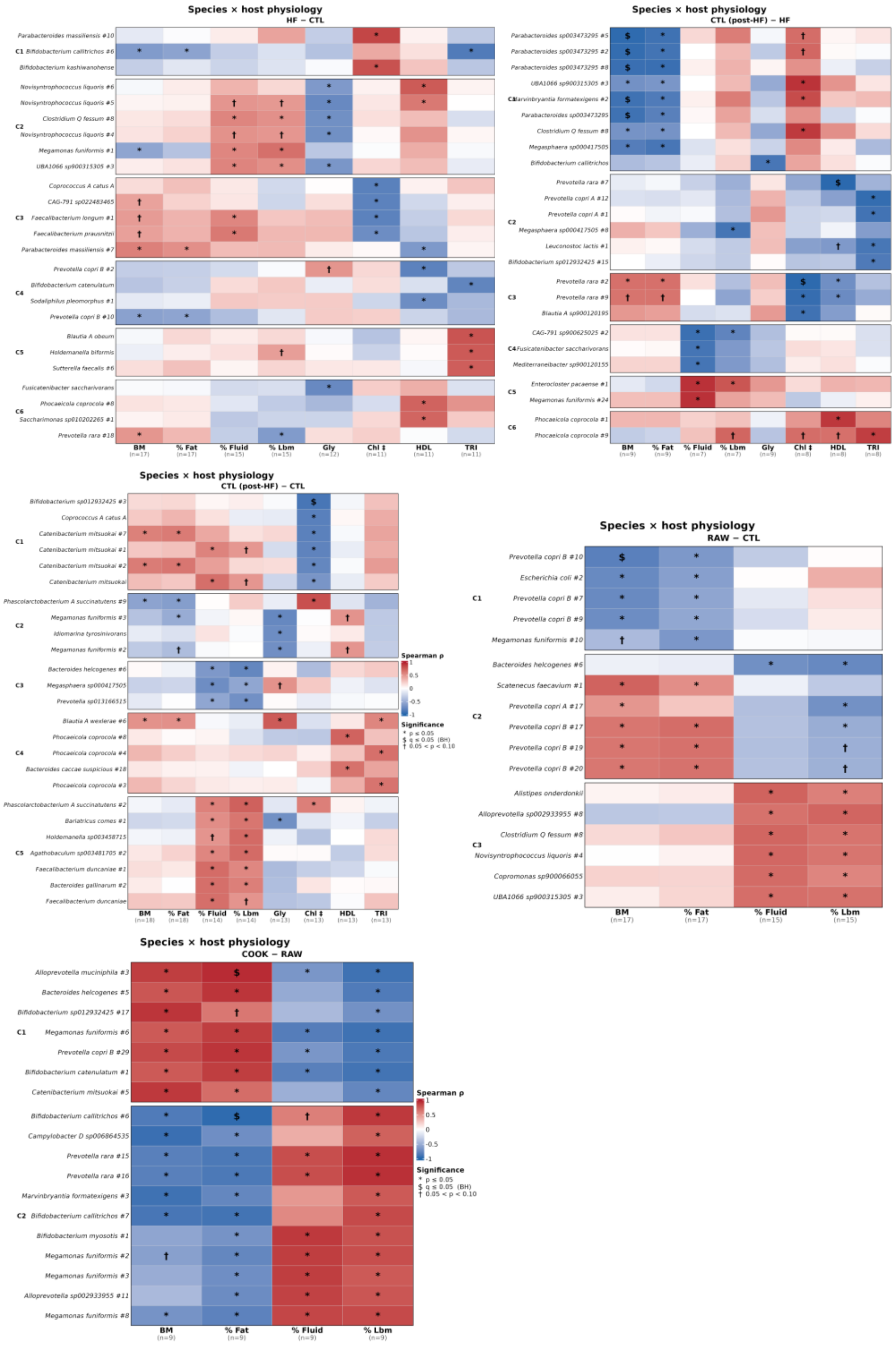


**Figure S4. Associations between gut-microbiome features and host metabolic and energetic phenotypes.** To Spearman rank correlations (ρ) between CLR-transformed species abundances and host phenotypes for the five diet contrasts, shown as one heatmap per contrast (top-five species per phenotype in rows, phenotypes in columns; fill: ρ from -1 in blue to +1 in red; * p ≤ 0.05, $ BH-adjusted q ≤ 0.05, † 0.05 < p < 0.10).
